# IFN*γ* and IFN*γ* mimetics prevent IFN-I-mediated TB susceptibility by regulating iron metabolism and lipid peroxidation

**DOI:** 10.64898/2026.07.01.735930

**Authors:** Prasanna Babu Araveti, Shivraj M. Yabaji, Muhammad Zainul Arifin, Suruchi Lata, Yuriy O. Alekseyev, Alexander A. Gimelbrant, William R. Bishai, Vadim Zhernovkov, Oleksii S. Rukhlenko, Boris N. Kholodenko, Igor Kramnik

## Abstract

Type I interferons (IFN-I) and IFNγ exert divergent effects during tuberculosis, but the mechanisms that determine whether macrophage activation promotes host defense or inflammatory pathology remain incompletely understood. Here, we dissect the interplay between IFN-I and IFNγ in macrophage activation using genetically susceptible B6.Sst1S macrophages. We show that, during tumor necrosis factor (TNF) stimulation, susceptible macrophages enter a persistent pathological activation state (pPAS) characterized by sustained lipid peroxidation and super-induction of IFN-I responses. This pathological state is maintained by autocrine IFN-I signaling. In contrast, IFNγ priming prevents pPAS development by enhancing macrophage resilience to oxidative stress, in part through regulation of iron metabolism and induction of ferritin expression. Computational **c**ell **s**tate **t**ransition **a**ssessment and **r**egulation (cSTAR) analysis identified pathways and small molecules predicted to promote the transition of susceptible macrophages toward an IFNγ-induced, Mtb-resistant state. Consistent with these predictions, the CDK4/6 inhibitor trilaciclib reduced lipid peroxidation by regulating iron metabolism, whereas retinoic acid signaling enhanced GPX4 expression and lipid biosynthesis programs. Combined CDK4/6 inhibition and retinoic acid receptor activation efficiently prevented the pathological activation state. Together, these findings delineate a mechanism of IFN-I/IFNγ crosstalk during macrophage activation and identify pharmacologic strategies to prevent IFN-I-dominant, lipid peroxidation-driven macrophage pathology.

## Introduction

Tuberculosis (TB), caused by *Mycobacterium tuberculosis* (Mtb), remains the leading cause of death by a single infectious agent worldwide, with 10.7 million new cases and 1.23 million deaths reported in 2024. Despite the availability of antimicrobial therapy, treatment failures occurred in 12% of TB patients (WHO, 2024), primarily attributed to the emergence of multidrug-resistant strains, HIV co-infections and host immune dysregulation (Furin et al., 2019; Pai et al., 2016). Remarkably, about a quarter of the global human population has been infected with Mtb, but only a fraction of those develop TB disease and transmits the bacteria, suggesting that the majority of immunocompetent humans are capable of spontaneously controlling and, possibly, eradicating Mtb (Behr et al., 2024). In contrast, the susceptible individuals, even immunocompetent, are not sufficiently protected by existing vaccines. Therefore, better understanding mechanisms of TB susceptibility are necessary to improve TB prevention and treatment outcomes in the susceptible individuals.

Among central unanswered questions are the mechanisms of dysregulation of macrophages that may either eradicate the bacteria or serve as the major host cell niche for the Mtb intracellular growth and persistence. Interferons are among the main known host factors that regulate macrophage interactions with virulent Mtb (Russell et al., 2025; Vance, 2025). Despite significant overlap of the transcriptional targets and signaling pathways of type I (IFN-I) and type II (interferon-gamma) interferons (Ravi Sundar Jose Geetha et al., 2024), their effects on TB are strikingly divergent: IFN-gamma (IFN*γ*) is essential for Mtb resistance, while IFN-I may induce a dysfunctional macrophage activation state associated with TB susceptibility (Moreira-Teixeira et al., 2018). IFN*γ* is produced by antigen-stimulated Th1 cells and natural killer (NK) cells and activates macrophages enabling them to control or kill the bacteria. It plays an essential role in host resistance to mycobacterial infections, including Mtb, as well as avirulent vaccine strain M. bovis BCG and opportunistic environmental mycobacteria in mice (Cooper et al., 1993; Flynn et al., 1993) and in humans (Fortin et al., 2007; Rosain et al., 2019). Among pleiotropic effects of IFN*γ* relevant to TB are the upregulation of macrophage effector functions, such as production of reactive oxygen and nitrogen species, phagocytosis, autophagy, phagosome maturation and targeting the bacteria for lysosomal degradation (Flynn et al., 1993; Gutierrez et al., 2004; MacMicking et al., 1997; Via et al., 1998). IFN*γ* promotes T cell-mediated immunity by inducing major histocompatibility complex (MHC) class II expression on the antigen-presenting cells and the formation of immunoproteasomes, which increase the peptide supply for the antigen presentation (O’Garra et al., 2013; Seifert et al., 2010). At the tissue and whole organism levels, IFN*γ* modulates hematopoiesis and immune cell trafficking (Kaufmann et al., 2018; MacNamara et al., 2011), regulates epithelial cell responses to injury (Li et al., 2024; Lin et al., 2024) and displays a paradoxical anti-inflammatory activity (Nandi & Behar, 2011).

The type I interferons (IFN-I) are mediators of innate immunity produced by multiple cells in response to viral infections, cytokines and stress (Boccuni et al., 2022). Compared to IFN*γ*, their roles in TB pathogenesis are more nuanced (Moreira-Teixeira et al., 2018; Munson & Kaushal, 2025). Thus, several experimental studies demonstrate protective effects of IFN-I pathway using macrophage infection with Mtb in vitro (Roberts et al., 2025) or during an early stage of Mtb infection in mice (Desvignes et al., 2012; Moreira-Teixeira et al., 2016). However, the overall impact of IFN-I on TB progression in vivo is negative. In human patients, transcriptomic studies identified an IFN-I inducible gene signature in the peripheral blood of active TB patients, which correlated with the disease severity and the risk of progression from latent to active disease (Berry et al., 2010; Cliff et al., 2013; Maertzdorf et al., 2011; Scriba et al., 2017). The upregulation of IFN-I pathways in human TB lesions was associated with TB severity (Fonseca et al., 2025). In animal TB models, more virulent Mtb strains induced higher levels of IFN-I that was associated with reduced Th1 immunity and decreased survival (Manca et al., 2005). Accordingly, inhibition of the IFN-I signaling using either the IFN-I receptor knockout or blocking antibodies reduced the inflammatory lung damage and increased the animal survival in several mouse models (Dorhoi et al., 2014; Ji et al., 2019; Moreira-Teixeira et al., 2020; Wiens & Ernst, 2016). Several mechanisms by which the IFN-I hyperactivity contributes to TB pathogenesis have been described. In vitro IFN-I suppressed macrophage responses to IFN*γ* (Teles et al., 2013) and exacerbated Mtb-induced macrophage death (Lee & Nathan, 2024). In vivo, IFN-I counters essential TB protective cytokine pathways: inhibits IL-12 and promotes the immunosuppressive IL-10 production (McNab et al., 2014); promotes macrophage polarization towards anti-inflammatory state (Benard et al., 2018); induces IL-1Ra, which blocks IL-1 (Ji et al., 2019), an essential protective cytokine (Mayer-Barber et al., 2010); promotes the neutrophil-driven lung damage (Chowdhury et al., 2024; Moreira-Teixeira et al., 2020; Saqib et al., 2025); suppresses T cell mobility and interactions with target cells in the lung (Branchett et al., 2025; Caouaille et al., 2025), and suppresses macrophage response to interferon gamma (Kotov et al., 2023; Teles et al., 2013; Zheng et al., 2025).

Our studies demonstrated a major role of the IFN-I pathway hyperactivity in the context of TB susceptible host – the *sst1*-susceptible mouse model of human-like pulmonary TB, which is characterized by the development of necrotic TB lesions (Kramnik et al., 2000; Pichugin et al., 2009; S. M. Yabaji, S. Lata, I. Gavrish, et al., 2025; S. M. Yabaji, S. Lata, A. E. Tseng, et al., 2025). In vitro the *sst1*-susceptible macrophages displayed an aberrant activation characterized by the superinduction of IFNβ that led to the development of the integrated stress response (Bhattacharya et al., 2021; He et al., 2013). The critical role of the IFN-I-driven ISR in vivo was further supported by the administration of the Integrated Stress Response Inhibitor (ISRIB) or blockade of the IFN-I pathway that reduced lung pathology and Mtb burdens in the *sst1*-susceptible congenic B6.Sst1S mice (Bhattacharya et al., 2021; Ji et al., 2019; Krug et al., 2023). Next, we demonstrated that the aberrant activation of the *sst1*-susceptible macrophages was primarily driven by coupling of the IFN-I pathway to dysfunctional antioxidant response, leading to the accumulation of lipid peroxidation products, unresolving stress and compromised Mtb resistance (Shivraj M. Yabaji et al., 2025).

Several lines of evidence suggest that boosting Th1-mediated adaptive immunity may correct the *sst1*-susceptible phenotype: subcutaneous immunization with BCG or Mtb delayed the pulmonary TB progression in the *sst1*-susceptible mice (Gern et al., 2025; Yan et al., 2007). It has been established that BCG vaccination induces Th1 response in the genetically resistant and susceptible mice (Kramnik et al., 1994) and in non-human primates (Simonson et al., 2025). The BCG-induced T lymphocytes restored the ability of the *sst1*-susceptible macrophages to control Mtb in vitro (Shivraj M. Yabaji et al., 2025). Therefore, we hypothesized that IFN*γ* may counter pathogenic effects of the IFN-I hyperactivity that drive TB progression. Moreover, an IFN*γ*-mimetic, rocaglamide A, identified in our previous studies, also prevented the aberrant macrophage activation and IFN*γ* superinduction (Bhattacharya et al., 2021; Chatterjee et al., 2021; Yabaji et al., 2023).

Here, we demonstrate that type I and type II interferons (IFN-I and IFN*γ*) antagonize each other in activated macrophages through differential regulation of iron metabolism and lipid peroxidation and propose small-molecule modulation of these pathways as a strategy to prevent pathogenic macrophage activation.

## Results

### 1. IFN-I autocrine signaling sustains pathological macrophage activation in a TNF-independent manner

The IFN-I and lipid peroxidation pathways underlying the aberrant activation of the *sst1*-susceptible macrophages by TNF are prone to auto-amplification. Therefore, we hypothesized that their coupling maintained this pathological activation state (PAS). To test this hypothesis, first we compared the activation states of the wild type B6 and the mutant B6.Sst1S bone marrow-derived macrophages (BMDMs) after TNF withdrawal. BMDMs were initially stimulated with TNF for 24 h, washed and cultured in fresh medium without exogenous TNF for an additional 16 h (Figure 1A). After this period, ROS production and *Ifnb1* mRNA expression in B6 BMDMs returned to baseline. In contrast, in B6.Sst1S macrophages we observed continuous ROS production, lipid peroxidation and *Ifnb1* mRNA upregulation (Figure 1B and 1C, Figure 1-figure supplement 1A, 1B). Consistent with the hyperactivity of the IFN-I pathway, we also observed the upregulation of *Rsad2* mRNA and downregulation of genes involved in lipid and cholesterol biosynthesis *Srebf2*, *Scd2*, and *Dhcr24* (Figure 1D, Figure 1-figure supplement 1C, 1D, and 1E) and unresponsiveness to IFN*γ* (Figure 1E). After blocking TNF signaling at 24 h, the level of intracellular 4-HNE adducts remained elevated exclusively in the B6.Sst1S macrophages (Figure 1-figure supplement 1J). Restimulation of B6.Sst1S macrophages with TNF at 24 h (Figure 1A) resulted in further upregulation of *Ifnb1* mRNA (Figure 1F, Figure 1-figure supplement 1F), increased levels of ROS and 4-HNE adducts (Figure 1G, Figure 1-figure supplement 1G and 1H) and persistent suppression of lipid biosynthesis genes (Figure 1H, Figure 1-figure supplement 1I). Thus, after initial TNF stimulation, the *sst1*-susceptible macrophages develop a self-sustained persistent pathological activation state (pPAS), which is characterized by continuous ROS production, dysregulated lipid metabolism with the accumulation of lipid peroxidation (LPO) products, unresponsiveness to IFN*γ* and an aberrant response to TNF restimulation, leading to further IFN-I pathway escalation.

**Figure 1:**
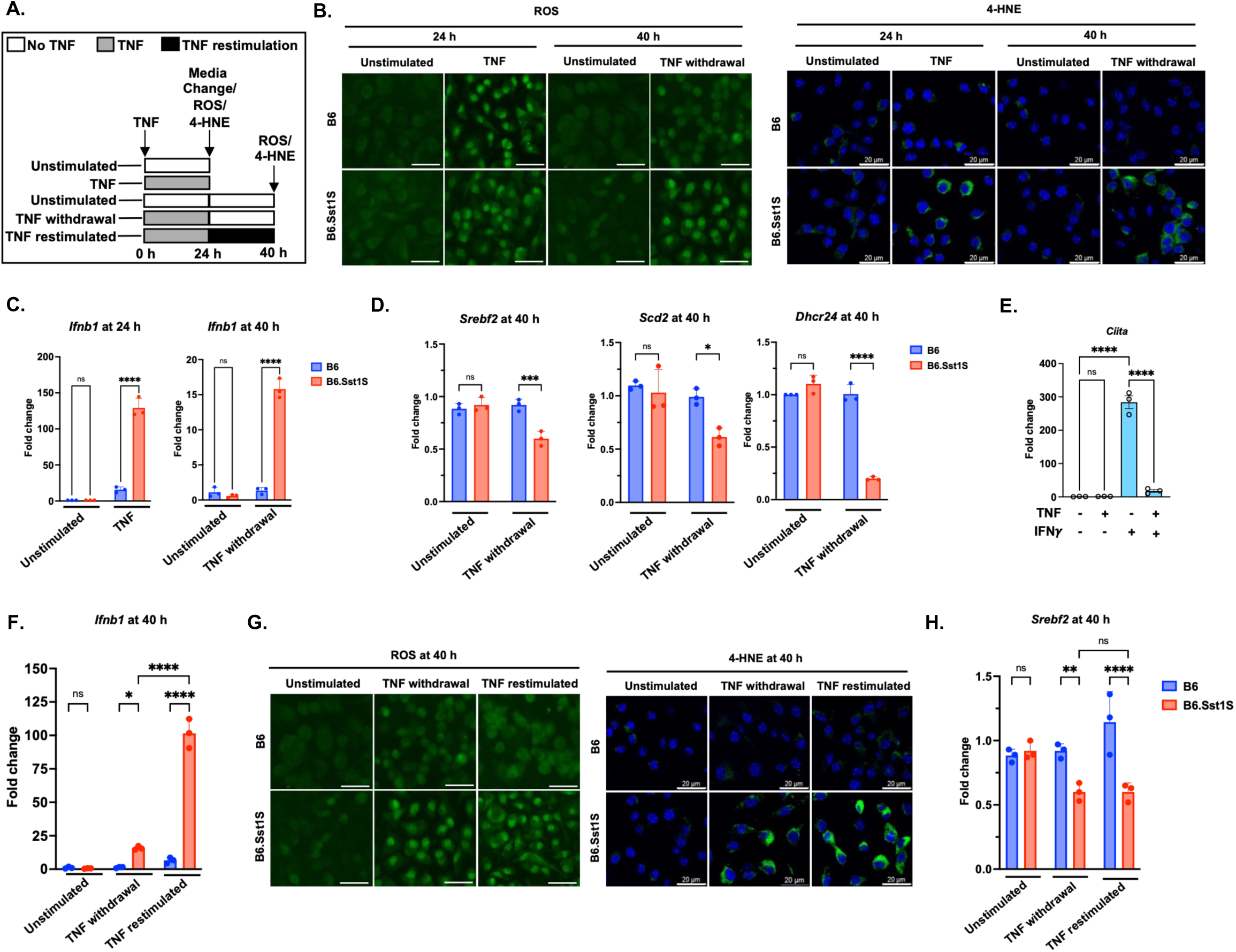
Susceptible B6.Sst1S macrophages undergo a persistent pathological activation state (pPAS) during TNF stimulation. A. Schematic of the experimental workflow. BMDMs from B6 and B6.Sst1S mice were stimulated with 10 ng/mL TNF for 24 h. Cells were either harvested immediately or cultured for an additional 16 h in fresh medium containing either TNF (TNF restimulation) or no TNF (TNF withdrawal) before harvest at 40 h after the initial stimulation. B. Persistent accumulation of ROS and 4-HNE adducts in B6.Sst1S BMDMs. BMDMs from B6 and B6.Sst1S mice were treated as described in panel A. Intracellular ROS levels (left panel) and accumulation of 4-HNE adducts (right panel) were subsequently assessed. ROS accumulation was detected using CellROX Green staining, and fluorescence images were acquired by fluorescence microscopy. Accumulation of the lipid peroxidation product 4-HNE was detected by confocal microscopy using a 4-HNE-specific antibody. Scale bar: 20 μm. C. IFN-I signaling remains persistently super-induced in B6.Sst1S BMDMs following TNF withdrawal. BMDMs from B6 and B6.Sst1S mice were treated as described in panel A. The mRNA expression levels of *Ifnb1* were quantified by qRT-PCR at 24 and 40 h post stimulation. Fold change was calculated relative to the unstimulated B6 control using the ΔΔCt method with 18S as an internal control. D. Lipid biosynthesis genes remain downregulated in B6.Sst1S BMDMs following TNF withdrawal. BMDMs from B6 and B6.Sst1S mice were treated as described in panel A. The mRNA expression levels of *Srebf2, Scd2*, and *Dhcr24* were quantified by qRT-PCR at 40 h post TNF stimulation. Fold change was calculated relative to the unstimulated B6 control using the ΔΔCt method with 18S as an internal control. E. B6.Sst1S BMDMs fail to respond to IFNγ during the pPAS state. B6.Sst1S BMDMs were stimulated with TNF for 18 h. After 18 h of TNF stimulation, IFNγ ( 5 U/mL) was added to the culture media, and cells were harvested at 40 h. The mRNA expression levels of *Ciita* was quantified by qRT-PCR. Fold change was calculated relative to the unstimulated control using the ΔΔCt method with 18S as an internal control. F. TNF restimulation during pPAS further enhances IFN-I signaling in B6.Sst1S BMDMs. BMDMs derived from B6 and B6.Sst1S mice were stimulated with TNF as described in panel A, including TNF restimulation at 24 h. Cells were harvested at 40 h, and *Ifnb1* mRNA expression levels were quantified by qRT-PCR. Fold change was calculated relative to the unstimulated B6 control using the ΔΔCt method with 18S as an internal control. G. TNF restimulation sustains elevated ROS levels and further increases 4-HNE adduct accumulation in B6.Sst1S BMDMs. BMDMs derived from B6 and B6.Sst1S mice were treated as described in panel A, including TNF restimulation at 24 h. Intracellular ROS levels were assessed using CellROX Green staining, and fluorescence images were acquired by fluorescence microscopy (left panel). Accumulation of 4-HNE adducts was detected by confocal microscopy using a 4-HNE-specific antibody (right panel). Scale bar: 20 μm. H. Lipid biosynthesis genes remain suppressed in B6.Sst1S BMDMs during TNF restimulation. BMDMs derived from B6 and B6.Sst1S mice were treated as described in panel A, including TNF restimulation at 24 h. The mRNA expression levels of *Srebf2* were quantified by qRT-PCR at 40 h. Fold change was calculated relative to the unstimulated B6 control using the ΔΔCt method with 18S as an internal control. The data are presented as mean ± standard deviation (SD) from three-five samples per experiment, representative of three independent experiments. The statistical analysis was performed by two-way ANOVA with Tukey’s multiple comparisons test (Panel D and E) and Sidak’s multiple comparison test (Panel F and H)), and one-way ANOVA with Tukey’s multiple comparisons test (Panel C) Significant differences are indicated with asterisks (ns, non significant; *, p < 0.05; **, p < 0.01; ***, p < 0.001; ****, p < 0.0001).

To test the role of IFN-I in the pPAS maintenance, we blocked IFN-I signaling using IFNAR1-specific antibodies after 24 h of TNF stimulation (Figure 2A). This blockade restored the B6.Sst1S macrophage responsiveness to IFN*γ* (Figure 2B) and prevented the 4-HNE adducts accumulation during the 24 – 40 h interval (Figure 2C, Figure 2-figure supplement 1A). The blockade of IFN-I signaling at the PAS initiation phase, i.e. at 2 h or 12 h after TNF stimulation, also significantly reduced the rates of lipid peroxidation (LAA biosynthesis) at 12 and 24 h of TNF stimulation, respectively (Figure 2D, Figure 2-figure supplement 1B and 1C). These data suggest that IFN-I signaling was a major driver of lipid peroxidation in TNF-activated B6.Sst1S macrophages at both the pPAS initiation and maintenance stages.

**Figure 2:**
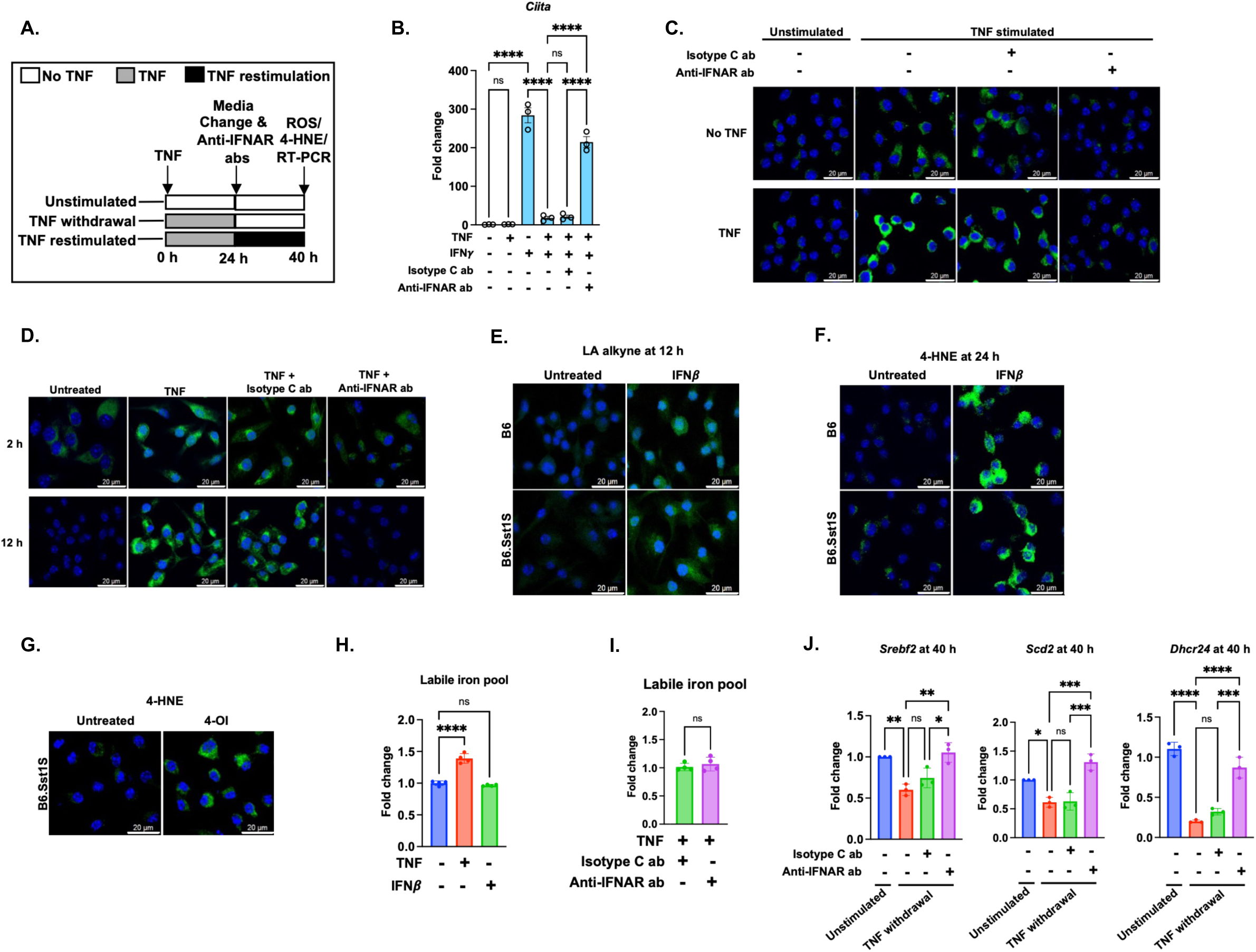
IFN-I autocrine signaling sustains pathological macrophage activation in TNF-independent manner. A. Schematic of the experimental workflow. BMDMs from B6 and B6.Sst1S mice were stimulated with 10 ng/mL TNF for 24 h. Cells were either harvested immediately or cultured for an additional 16 h following replacement with fresh medium containing either TNF (TNF restimulation) or no TNF (TNF withdrawal). Where indicated, cells additionally received isotype control antibodies or anti-IFNAR antibodies (5 μg/mL) at the time of media replacement. Cells were harvested at 40 h after the initial TNF stimulation. B. IFN-I signaling suppresses IFNã responsiveness in B6.Sst1S BMDMs during the pPAS state. B6.Sst1S BMDMs were stimulated with TNF for 18 h, followed by addition of IFNã (5 U/mL) in the presence of either isotype control antibodies or anti-IFNAR antibodies, where indicated. Cells were harvested at 40 h after the initial TNF stimulation, and *Ciita* mRNA expression was quantified by qRT-PCR. Gene expression was normalized to 18S and expressed relative to unstimulated controls using the ΔΔCt method. C. IFN-I sustains the 4-HNE adducts accumulation in B6.Sst1S BMDMs during pPAS. BMDMs from B6.Sst1S mice were treated as described in A. The accumulation of lipid peroxidation product, 4-HNE was detected by confocal microscopy using 4-HNE specific antibody. Scale bar-20 *μ*m. D. IFN-I signaling initiates and sustains lipid peroxidation product synthesis in B6.Sst1S BMDMs during TNF stimulation. B6.Sst1S BMDMs were stimulated with TNF, and either isotype control antibodies or anti-IFNAR antibodies were added at 2 h or 12 h after TNF stimulation. Cells receiving antibody treatment at 2 h were harvested at 12 h following TNF stimulation (top panel), whereas cells receiving antibody treatment at 12 h were harvested at 24 h following TNF stimulation (bottom panel). Lipid peroxidation product synthesis was assessed by LA alkyne staining. Representative fluorescence images are shown. Scale bar, 20 μm. E. IFNâ induces lipid peroxidation product synthesis independently of the *sst1* locus. BMDMs from B6 and B6.Sst1S mice were treated with IFNβ (300 U/mL) for 12 h, followed by assessment of lipid peroxidation product synthesis by LA alkyne staining. Representative fluorescence images are shown. Scale bar, 20 μm. F. IFNâ induces 4-HNE adduct accumulation independently of the *sst1* locus. BMDMs from B6 and B6.Sst1S mice were treated with IFNβ (300 U/mL) for 24 h, followed by immunofluorescence staining for 4-HNE adducts using 4-HNE specific antibodies. Representative confocal images are shown. Scale bar, 20 μm. G. Direct 4-octyl itaconate treatment induces 4-HNE adduct accumulation in B6.Sst1S BMDMs. B6.Sst1S BMDMs were treated with 4-octyl itaconate (4-OI) (5 *μ*M) for 24 h, followed by immunofluorescence staining for 4-HNE adducts using 4-HNE specific antibodies. Representative confocal images are shown. Scale bar, 20 μm. H. IFNâ does not alter the labile iron pool in B6.Sst1S BMDMs. B6.Sst1S BMDMs were treated with TNF (10 ng/mL) or IFNβ (300 U/mL) for 24 h and the cells were stained with FerroOrange to assess intracellular labile iron levels. Representative fluorescence images are shown. Fluorescence intensity was quantified and expressed as fold change relative to untreated controls. Scale bar, 20 μm. I. IFN-I signaling does not regulate iron metabolism during TNF stimulation in B6.Sst1S BMDMs. B6.Sst1S BMDMs were stimulated with TNF (10 ng/mL), followed by addition of either isotype control antibodies or anti-IFNAR antibodies at 2 h post stimulation. Cells were stained with FerroOrange at 24 h to assess intracellular labile iron levels. Fluorescence intensity was quantified and expressed as fold change relative to isotype control antibody-treated samples. J. IFN-I signaling suppresses lipid biosynthesis gene expression during the pPAS state. B6.Sst1S BMDMs were treated as described in panel A under the TNF withdrawal condition. Cells were harvested at 40 h after the initial TNF stimulation, and expression of Srebf2, Scd2, and Dhcr24 was quantified by qRT-PCR. Gene expression was normalized to 18S and expressed relative to unstimulated controls using the ΔΔCt method. The data are presented as mean ± SD from three-five samples per experiment, representative of three independent experiments. The statistical analysis was performed by two-way ANOVA with Tukey’s multiple comparisons test (Panel J), and one-way ANOVA with Tukey’s multiple comparisons test (Panel B, H, and I). Significant differences are indicated with asterisks (ns, non significant; *, p < 0.05; **, p < 0.01; ***, p < 0.001; ****, p < 0.0001).

Accordingly, treatment of naïve B6 and B6.Sst1S BMDMs with IFN*β* alone induced lipid peroxidation, as determined by the rate of LAA production (Figure 2E) and the intracellular 4-HNE adducts accumulation (Figure 2F), which occurred in a *sst1*-independent manner. We hypothesized that this effect could be mediated by itaconate, an electrophilic metabolite produced by the IFN-inducible enzyme Acod1 (Lampropoulou et al., 2016; Peace & O’Neill, 2022). Indeed, treatment of naïve BMDMs with 4-octyl itaconate (4-OI) led to the 4-HNE adducts accumulation (Figure 2G, Figure 2-figure supplement 1D).

Considering other factors involved in lipid peroxidation in TNF-stimulated macrophages, IFN-I signaling neither affected the levels of the intracellular labile iron pool (Figure 2H and 2I, respectively), nor did it appeared to be a major driver of ROS production (Figure 2-figure supplement 1F and 1G). In these settings, however, IFN-I suppressed lipid biosynthesis genes, as evidenced by their upregulation after the IFN-I receptor blockade (Figure 2J, Figure 2-figure supplement 1E).

Taken together, these data suggest that persistent IFN-I signaling maintains pPAS by stimulating itaconate biosynthesis and suppressing lipid biosynthesis genes, thus promoting the accumulation of toxic lipid peroxidation products.

### 2. IFNγ priming prevents pPAS by limiting intracellular iron availability

Previous studies demonstrated that the induction of Th1-mediated immunity partially corrected the *sst1*-mediated Mtb susceptibility in vivo (Gern et al., 2025). Accordingly, co-stimulation of the B6.Sst1S BMDMs with IFN*γ* and TNF significantly increased their ability to control the intracellular Mtb growth in vitro (Figure 3A). Therefore, we tested whether macrophage priming with IFN*γ* could prevent or ameliorate pPAS induced by TNF in the susceptible macrophages (Figure 3-figure supplement 1A).

**Figure 3:**
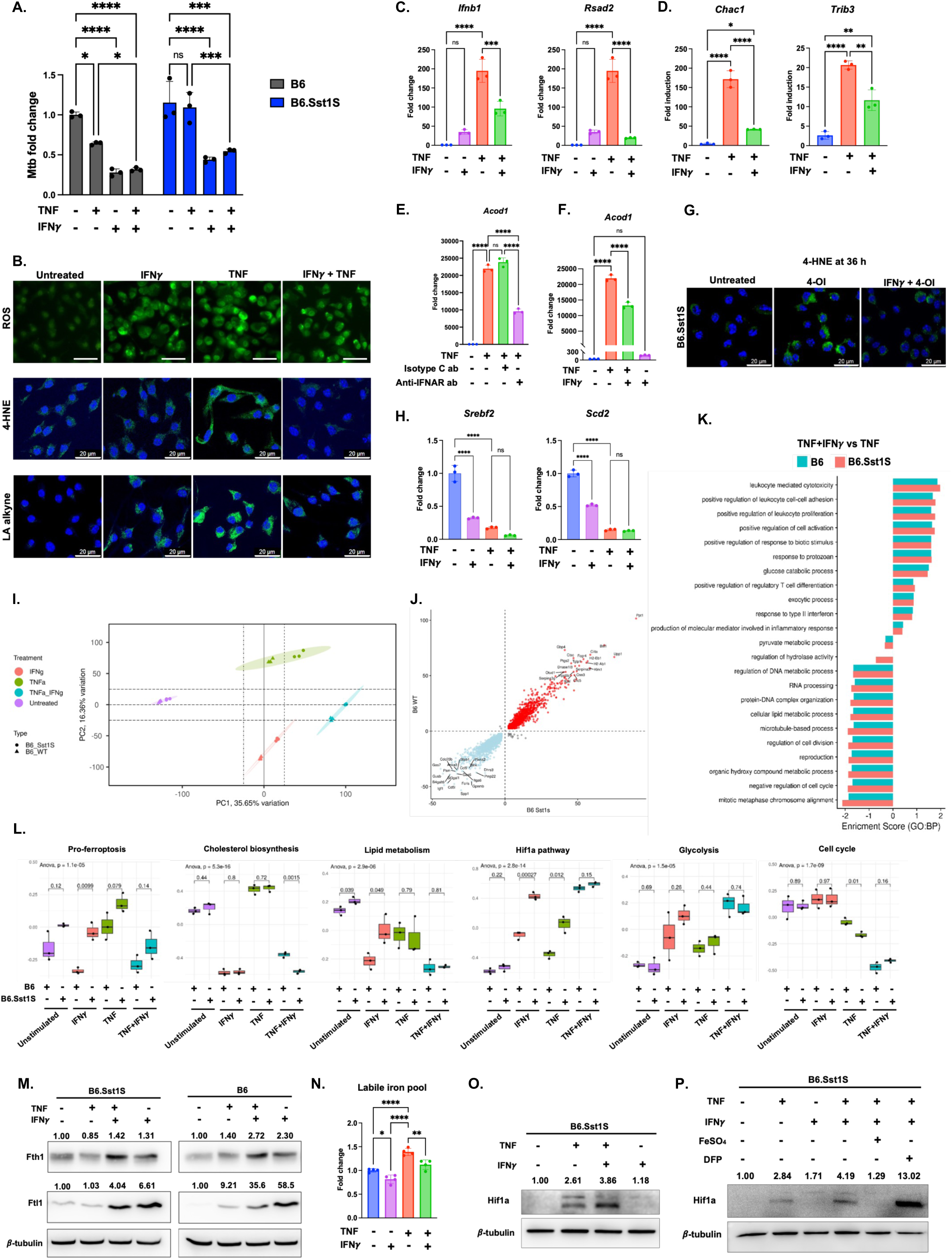
IFN*γ* priming prevents pPAS. A. IFNã controls intracellular Mtb growth in both B6 and B6.Sst1S BMDMs following TNF stimulation. BMDMs were stimulated with TNF (10 ng/mL) in the presence or absence of IFNγ (5 U/mL). The following day, cells were infected with Mtb Erdman reporter strain (SSB-GFP, smyc’::mCherry) at MOI = 1 and cultured for 5 days. On day 5 post-infection, intracellular Mtb load was quantified by qRT-PCR-based method. B. IFNã priming attenuates TNF-induced oxidative stress and lipid peroxidation in B6.Sst1S BMDMs. B6.Sst1S BMDMs were primed with IFNã (5 U/mL) for 16 h, followed by TNF (10 ng/mL) stimulation for 36 h (24 h for LA alkyne). ROS levels were assessed using CellROX Green staining and quantified by fluorescence microscopy (top panel). Accumulation of 4-HNE adducts was evaluated by confocal microscopy using a 4-HNE-specific antibody (middle panel). Lipid peroxidation product synthesis was assessed by LA alkyne staining, with images acquired by confocal microscopy (bottom panel). Representative images are shown. Scale bar, 20 μm. C-D. IFN*γ* priming prevents IFN-I super-induction and integrated stress response during TNF stimulation. B6.Sst1S BMDMs were primed with IFNγ (5 U/mL) for 16 h, followed by TNF (10 ng/mL) stimulation for 24 h. The mRNA levels of *Ifnb1* and *Rsad2* (Panel C) and *Trib3* and *Chac1* (Panel D) were quantified by qRT-PCR. The fold change was calculated normalizing with the untreated control using ΔΔCt method using 18S as an internal control. E. IFN-I maintains *Acod1* overexpression during TNF stimulation in B6.Sst1S BMDMs. B6.Sst1S BMDMS were treated with TNF (10 ng/mL) for 40 h and the mRNA levels of *Acod1* was quantified by qRT-PCR. The fold change was calculated normalizing with the untreated control using ΔΔCt method using 18S as an internal control. Isotype control or anti-IFNAR antibodies were added to the culture at 2 h after TNF stimulation. F. IFNã priming prevents *Acod1* overexpression during TNF stimulation. B6.Sst1S BMDMs were either primed with IFNγ (5 U/mL) for 16 h or left unprimed, followed by treatment with TNF (10 ng/mL) for an additional 24 h. The *Acod1* mRNA expression was quantified by qRT-PCR and presented as fold change. Gene expression was normalized to 18S and expressed compared to unstimulated controls using the ΔΔCt method. G. IFN*γ* priming prevents 4-OI induced 4-HNE adducts accumulation. B6.Sst1S BMDMs were either primed with IFNγ (5 U/mL) for 16 h or left unprimed, followed by treatment with 4-octyl itaconate (4-OI; 5 *μ*M) for an additional 36 h. 4-HNE adducts accumulation was observed by confocal microscopy using 4-HNE specific antibody. Scale bar-20 *μ*m. H. IFN*γ* priming do not restore lipid biosynthesis gene expression during TNF stimulation. B6.Sst1S BMDMs were primed with IFNγ (5 U/mL) for 16 h, followed by TNF (10 ng/mL) stimulation for an additional 24 h. The mRNA levels of *Srebf2* and *Scd2* were quantified by qRT-PCR and presented as fold change. Gene expression was normalized to 18S and expressed compared to unstimulated controls using the ΔΔCt method. I. The PCA plot showing the combination of IFN*γ* and TNF leads to a distinct transcriptomic profile compared to the individual counterparts with least variation between B6 and B6.Sst1S macrophages. B6 and B6.Sst1S BMDMs were treated individually with IFN*γ* (5 U/mL) or TNF (10 ng/mL) or in combination for 12 h. Cells were harvested, and transcriptome was analyzed by RNA-seq. J. The combination of IFN*γ* and TNF causes similar transcriptomic profile in B6 and B6.Sst1S macrophages. The differential gene expression analysis showing the significantly upregulated and downregulated genes were common in B6 and B6.Sst1S macrophages when treated in combination of IFN*γ* and TNF compared to TNF alone. The differentially expressed genes with adjusted p-value < 0.05 and absolute fold change >1.5 were considered. K. The type II interferon response, glucose catabolic processes were upregulated, and cell cycle related processes were downregulated in both B6 and B6.Sst1S macrophages upon stimulation in combination of IFN*γ* and TNF compared to TNF alone. The gene set enrichment analysis based on biological processes comparing the IFN*γ* and TNF combination treatment to TNF alone in B6 and B6.Sst1S macrophages. The significant enriched pathways were defined with adjusted p-value < 0.05. L. The single sample gene set enrichment analysis showing downregulation of genes involved in pro-ferroptosis, cholesterol biosynthesis, lipid metabolism, cell cycle and upregulation of genes involved in Hif1a pathway, glycolysis when treated with both IFN*γ* and TNF in B6 and B6.Sst1S macrophages independent of sst1. M. IFN*γ* priming enhances ferritins heavy chain and ferritin light chain expression independent of sst1. B6.Sst1S BMDMs were primed with IFNγ (5 U/mL) for 16 h, followed by TNF (10 ng/mL) stimulation for 24 h. and Fth1 and Ftl1 protein levels were quantified by Western immunoblotting. *β*-tubulin was used as loading control. Average densitometric values from two independent experiments were included above the blot. N. IFN*γ* priming suppresses labile iron pool during TNF stimulation. B6.Sst1S BMDMs were primed with IFNγ (5 U/mL) for 16 h, followed by TNF (10 ng/mL) stimulation for 24. After 24 h the cells were stained with FerroOrange and fluorescence intensities were quantified. Bar graphs showing the fold change in fluorescence intensities compared to the untreated samples. (n=4). O. IFN*γ* priming enhances Hif1a protein levels in B6.Sst1S BMDMs. B6.Sst1S BMDMs were primed with IFNγ (5 U/mL) for 16 h, followed by TNF (10 ng/mL) stimulation for 24 h and HIF1*α* protein levels were quantified by Western immunoblotting. *β*-tubulin was used as loading control. Average densitometric values from two independent experiments were included above the blot. P. The iron metabolism regulated by IFN*γ* priming enhances Hif-1*α* protein levels during TNF stimulation in B6.Sst1S BMDMs. B6.Sst1S BMDMs were primed with IFNγ (5 U/mL) for 16 h, followed by stimulation with TNF (10 ng/mL) alone or in presence of either FeSO4 or deferiprone (DFP) for 24 h. Hif1a protein levels were quantified by Western immunoblotting. *β*-tubulin was used as loading control. Average densitometric values from two independent experiments were included above the blot. The data are presented as mean ± SD from three-five samples per experiment, representative of three independent experiments. The statistical analysis was performed by two-way ANOVA with Tukey’s multiple comparisons test (Panel A), and one-way ANOVA with Tukey’s multiple comparisons test (Panel C, D, E, F,H, and N). Significant differences are indicated with asterisks (ns, non significant; *, p < 0.05; **, p < 0.01; ***, p < 0.001; ****, p < 0.0001).

Although IFN*γ* alone moderately stimulated ROS production and LPO synthesis, the IFN*γ* priming significantly reduced LPO synthesis and accumulation induced by TNF (Figure 3B, Figure 3-figure supplement 1B). Moreover, IFN*γ* priming prevented the *Ifnb1* super-induction (Figure 3C), and reduced the upregulation of IFN-I-driven pPAS markers, such as *Rsad2*, the integrated stress response (ISR) genes, *Trib3* and *Chac1*, and *Acod1* (Figure 3C, 3D, 3F). Of note, IFN*γ* priming also suppressed the 4-HNE adduct accumulation caused by synthetic itaconate 4-OI (Figure 3G, Figure 3-figure supplement 1C). The latter observations suggested that IFN*γ* priming prevented the LPO accumulation independently of the IFN-I - Acod1 – itaconate downregulation. In addition, the IFN*γ* priming suppressed the expression of lipid biosynthesis genes, similarly to the IFN-I mediated effect of TNF described above (Figure 3H, Figure 3-figure supplement 1D). Also, IFN*γ* priming did not enhance the GPX4 expression in B6.Sss1S BMDMs (Figure 3-figure supplement 1E), which is maintained at lower levels compared to the B6 BMDMs during TNF stimulation (Shivraj M. Yabaji et al., 2025). The latter observations suggested that IFN*γ* prevented the LPO accumulation independently of the downregulation of the IFN-I - Acod1 – itaconate axis and lipid biosynthesis.

To reveal the preventive effect of IFN*γ* on pPAS development, we compared transcriptional responses of the B6 and B6.Sss1S BMDMs to IFN*γ*, TNF and their combination at the pPAS incipient stage, i.e. at 12 h after the cytokine stimulation, using bulk RNA-seq. The co-stimulation with IFN*γ* and TNF induced similar shifts in cell states and reduced the PC1 distance between the resistant and susceptible macrophages, as evidenced by the Principal Component Analysis (PCA) (Figure 3I), and by the similarities of their transcriptomes (Figure 3J). Thus, the MHC class II genes were upregulated, and the genes related to tissue remodeling, alternative macrophage activation, and cell division were downregulated in both B6 and B6.Sst1S macrophages (Figure 3J). The pathway analysis and gene set variation analysis (GSVA) also demonstrated similar modulation of TNF responses by IFN*γ* in BMDMs of both genetic backgrounds (Figure 3K and 3L). Among key metabolic pathways, the Hypoxia-Inducible Factor 1-alpha (Hif1a) and glycolysis pathways were upregulated, while the cell cycle, cholesterol biosynthesis and lipid metabolism pathways were downregulated in agreement with other published studies. Importantly, we found that IFN*γ* co-stimulation downregulated the expression of pro-ferroptic gene set in TNF-stimulated B6.Sst1S macrophages to the B6 level (Figure 3L).

As we reported previously, an impairment of the anti-oxidant response in TNF-stimulated B6.Sst1S macrophages, led to the LPO accumulation. In contrast to B6, the B6.Sst1S macrophages failed to upregulate ferritin light and heavy chains upon TNF stimulation and, thus, did not reduce the intracellular labile iron pool that sustained lipid peroxidation (Shivraj M. Yabaji et al., 2025). In contrast, IFN*γ* upregulated Fth1 and Ftl1 proteins in the s*st1*-independent manner either alone or in synergy with TNF (Figure 3M) and significantly reduced the labile iron pool in the B6.Sst1S macrophages (Figure 3N). We propose that this IFN*γ* effect plays a major role in preventing LPO accumulation in macrophages maximally activated by the IFN*γ* and TNF co-stimulation and, therefore, producing copious amounts of ROS.

In agreement with the Hif1a pathway upregulation detected by GSVA, we demonstrated upregulation of the Hif1a protein level in IFN*γ* and TNF co-stimulated macrophages, as compared to macrophages stimulated with TNF alone (Figure 3O). Because treatment with IFN*γ* alone did not upregulate the Hif1a protein level, we hypothesized that the TNF and IFN*γ* synergy could be explained by the known role of iron in regulating degradation of Hif1a (Knowles et al., 2006). Indeed, adding free iron to B6.Sst1S macrophages co-stimulated with IFN*γ* and TNF abolished the Hif1a upregulation, while iron chelation using deferiprone (DFP) further increased the protein level (Figure 3P). Taken together, these data demonstrate that by reducing the intracellular labile iron pool, IFN*γ* prevents lipid peroxidation and promotes Hif1a-mediated metabolic reprogramming of macrophages, thereby increasing their resilience to oxidative stress and preventing self-sustained pathological activation.

### 3. IFNγ mimetics prevent pPAS

To determine how to mimic the effect of IFN*γ* using small molecules, we used cell State Transition Assessment and Regulation (cSTAR) analysis, a recently developed systems biology approach that integrates multi-omics data to map distinct cell states and predict perturbations controlling transitions between them (Kholodenko et al., 2023; Rukhlenko et al., 2022).

We used RNA-sequencing (RNA-seq) data to map the cytokine priming states of B6 and B6.Sst1S macrophages following a 12 h stimulation with TNF, IFN*γ*, and their combination (Figure 4A). To capture these dynamics, we built a State Transition Vector (STV_TB_). Unlike previous works relying on classification methods (Rukhlenko et al., 2022; Yabaji et al., 2023), we utilized regression methods to achieve higher quantitative precision. This STV_TB_ determines the direction of gene expression changes towards the most TB-resistant state while integrating both genetic background (B6 vs. B6.Sst1S) and cytokine priming effects. Dynamic phenotype descriptor (DPD_TB_) scores were then calculated to quantify the phenotypic shifts induced by each perturbation (Rukhlenko et al., 2022; Yabaji et al., 2023). It can be seen (Figure 4B) that TNF stimulation of B6.Sst1S macrophages decreased their DPD_TB_ score, in agreement with their pathological activation state. In contrast, TNF and IFN*γ* stimulation of B6 macrophages increased their DPD_TB_ scores, with the highest score obtained by combining both cytokines, reflecting their known synergistic effect on macrophage activation. Although IFN*γ* and IFN*γ* + TNF stimulation of B6.Sst1S macrophages resulted in increased DPD_TB_ scores, these responses were significantly weaker than those observed in B6 macrophages (Figure 4A). The calculated DPD_TB_ scores strongly correlated with Mtb fold change data (Figure 4B). Thus, in these settings, a negative DPD_TB_ is indicative of Mtb-permissive macrophage cell state, whereas an increase in DPD_TB_ score is indicative of increased Mtb resistance.

**Figure 4:**
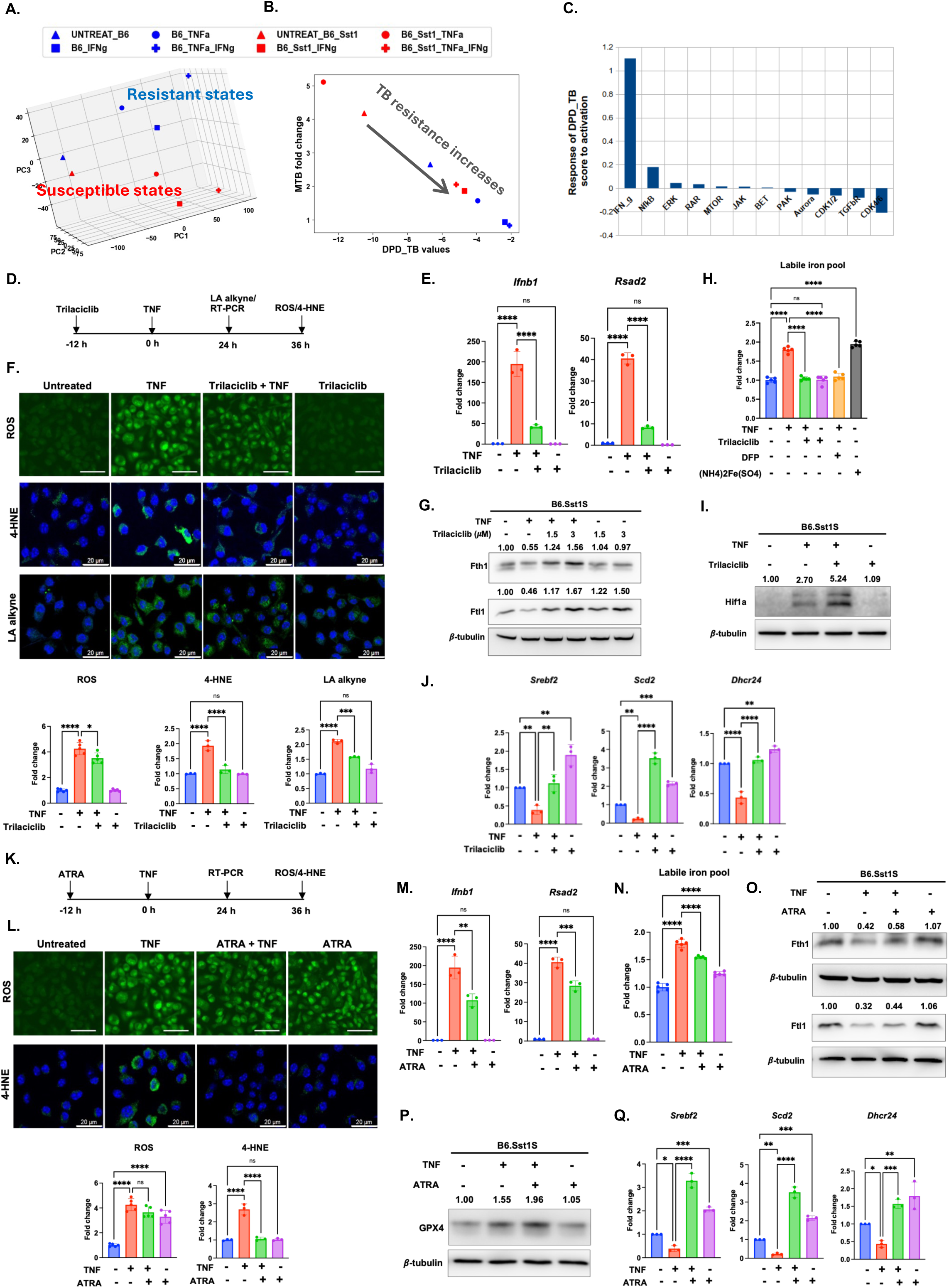
IFN*γ* mimetics prevents pPAS. A. The PCA plot illustrating qualitative separation between TB-resistant and TB-sensitive states. Each dots shows average of 3 biological replicates, presented individually at Figure 2I. B. Comparison of DPD_TB scores of macrophage resistance towards Mtb (DPD_TB) calculated for B6.Sst1S (red) and B6 (blue) macrophages treated with TNF (10 ng/mL), IFN*γ* (5 U/mL) and their combination for 12 hours. The DPD scores were computed using the changes in macrophage transcriptomics patterns. Higher DPD_TB scores indicate the greater the Mtb resistance. C. cSTAR network predicted responses of the DPD_TB phenotypic score to activation of core network modules. The responses of the DPD_TB score were calculated using the inferred influences of the core network pathways depicted in (C). Negative values represent decreases in macrophage resistance to Mtb, positive values indicate enhanced Mtb resistance following an increase in each core pathway activity. D. Schematic of the experimental workflow. B6.Sst1S BMDMs were pretreated with CDK4/6 inhibitor, trilaciclib (3 *μ*M) for 12 h followed by TNF (10 ng/mL) stimulation for 24 h or 36 h. E. Trilaciclib pretreatment prevents IFN-I super-induction during TNF stimulation. B6.Sst1S BMDMs were pretreated with trilaciclib (3 *μ*M) for 12 h followed by TNF stimulation for additional 24 h. The mRNA levels of *Ifnb1* and *Rsad2* were quantified by qRT-PCR. The fold change was calculated by normalizing with the untreated control using ΔΔCt method. 18S was used an an internal control. F. Trilaciclib pretreatment reduces ROS accumulation and lipid peroxidation during TNF stimulation. B6.Sst1S BMDMs were pretreated with trilaciclib (3 *μ*M) for 12 h, followed by TNF (10 ng/ml) stimulation for an additional 36 h (24 h for LA alkyne staining). Separate control groups were treated with either TNF alone or trilaciclib alone. Intracellular ROS levels were assessed using CellROX Green staining and quantified by fluorescence microscopy (top panel). Accumulation of 4-HNE adducts was evaluated by confocal microscopy using a 4-HNE-specific antibody (middle panel). Lipid peroxidation product synthesis was assessed by LA alkyne staining, and images were acquired by confocal microscopy (bottom panel). Representative images are shown. Scale bar: 20 μm. Bar graphs show the fold change in fluorescence intensity relative to untreated controls. Data represent mean ± SEM from three independent experiments (n=3). G. Trilaciclib pretreatment enhances ferritins heavy chain and ferritin light chain expression during TNF stimulation. B6.Sst1S BMDMs were pretreated with trilaciclib for 12 h followed by TNF stimulation for additional 24 h. Fth1 and Ftl1 protein levels were measured by western immunoblotting. Average densitometric values from two independent experiments were included above the blot. *β*-tubulin was used as loading control. H. Trilaciclib pretreatment suppresses labile iron pool during TNF stimulation. B6.Sst1S BMDMs were pretreated with trilaciclib (3 *μ*M) for 12 h followed by TNF (10 ng/mL) stimulation for additional 24 h. Cells were stained with FerroOrange, and fluorescence intensities were quantified by florescence microscopy. Bar graphs showing the fold change in fluorescence intensities compared to the untreated samples. (n=5). I. Trilaciclib pretreatment enhances Hif1a protein levels in B6.Sst1S BMDMs. B6.Sst1S BMDMs were pretreated with trilaciclib (3 *μ*M) for 12 h followed by TNF stimulation for additional 24 h. Hif1a protein level was measured by Western immunoblotting. Average densitometric values from two independent experiments were included above the blot. *β*-tubulin was used as loading control. J. Trilaciclib pretreatment enhances lipid biosynthesis gene expression during TNF stimulation. B6.Sst1S BMDMs were pretreated with trilaciclib (3 *μ*M) for 12 h followed by TNF stimulation for additional 24 h. The mRNA levels of *Srebf2*, *Scd2*, and *Dhcr24* were quantified by qRT-PCR. The fold change was calculated normalizing with the untreated control using ΔΔCt method. 18S was used as an internal control. K. Schematic of the experimental workflow. B6.Sst1S BMDMs were pretreated with All trans Retinoic acid (ATRA) (2 *μ*M) for 12 h followed by TNF (10 ng/mL) stimulation for 24 h or 36 h. L. ATRA pretreatment show minimal effect ROS levels during TNF stimulation. B6.Sst1S BMDMs were pretreated with ATRA (2 *μ*M) for 12 h, followed by TNF (10 ng/ml) stimulation for an additional 36 h. Separate control groups were treated with either TNF alone or ATRA alone. Intracellular ROS levels were assessed using CellROX Green staining and quantified by fluorescence microscopy (top panel). Accumulation of 4-HNE adducts was evaluated by confocal microscopy using a 4-HNE-specific antibody (bottom panel). Scale bar: 20 μm. Bar graphs show the fold change in fluorescence intensity relative to untreated controls. Data represent mean ± SEM from three independent experiments (n=3). M. ATRA pretreatment controls IFN-I super-induction during TNF stimulation. B6.Sst1S BMDMs were pretreated with ATRA (2 *μ*M) for 12 h followed by TNF stimulation for additional 24 h. The mRNA levels of *Ifnb1* and *Rsad2* were quantified by qRT-PCR. The fold change was calculated by normalizing with the untreated control using ΔΔCt method. 18S was used an an internal control. N. ATRA pretreatment reduces labile iron pool during TNF stimulation. B6.Sst1S BMDMs were pretreated with ATRA (2 *μ*M) for 12 h followed by TNF (10 ng/mL) stimulation for additional 24 h. Cells were stained with FerroOrange, and fluorescence intensities were quantified by florescence microscopy. Bar graphs showing the fold change in fluorescence intensities compared to the untreated samples. (n=5). O. ATRA pretreatment enhances ferritins heavy chain and ferritin light chain expression during TNF stimulation. B6.Sst1S BMDMs were pretreated with ATRA (2 *μ*M) for 12 h followed by TNF stimulation for additional 24 h. Fth1 and Ftl1 protein levels were measured by western immunoblotting. Average densitometric values from two independent experiments were included above the blot. *β*-tubulin was used as loading control. P. ATRA pretreatment enhances GPX4 protein expression in B6.Sst1S BMDMs. B6.Sst1S BMDMs were pretreated with ATRA (2 *μ*M) for 12 h followed by TNF stimulation for additional 24 h. GPX4 protein levels was measured by Western immunoblotting. Average densitometric values from two independent experiments were included above the blot. *β*-tubulin was used as loading control. Q. ATRA pretreatment enhances lipid biosynthesis gene expression during TNF stimulation. B6.Sst1S BMDMs were pretreated with ATRA (2 *μ*M) for 12 h followed by TNF stimulation for additional 24 h. The mRNA levels of *Srebf2*, *Scd2*, and *Dhcr24* were quantified by qRT-PCR. The fold change was calculated normalizing with the untreated control using ΔΔCt method. 18S was used as an internal control. The data are presented as mean ± SD from three-five samples per experiment, representative of three independent experiments. The statistical analysis was performed one-way ANOVA with Tukey’s multiple comparisons test (Panel E, F, H, J, L, M, N, and Q). Significant differences are indicated with asterisks (ns, non significant; *, p < 0.05; **, p < 0.01; ***, p < 0.001; ****, p < 0.0001).

By analyzing transcriptional responses to perturbations using publicly available perturbation transcriptomics datasets, cSTAR enables reconstruction of the protein signaling network wiring (Deng et al., 2025). Here, we applied cSTAR to reconstruct the network that controls the transition between the susceptible and resistant macrophage activation states and identified key signaling pathways (nodes) that govern macrophage priming for Mtb resistance. Predictions of how activation of each node by 1% will change DPD_TB_ score are shown in Figure 4C. As anticipated, the IFN*γ* pathway had the strongest positive effect on the DPD_TB_ score. Also, activation of NFkB, ERK, retinoic acid receptor (RAR), JAK/STAT and mTOR pathways contributed to the DPD_TB_ increase, i.e. predicted to move macrophage activation state towards higher Mtb resistance. In contrast, cyclin dependent kinases (CDK 4/6 and CDK1/2) and TGFβ receptor-mediated pathways reduced DPD_TB_, i.e. were predicted to compromise the resistant state. In summary, our cSTAR analysis predicted that stimulating the *sst1*-susceptible macrophages with TNF in combination with either CDK inhibition or RAR activation could partially mimic the IFN*γ* priming and enhance their Mtb resistance. Next, we tested these predictions experimentally.

To investigate the effect of CDK4/6 inhibition, we pre-treated B6.Sst1S macrophages with CDK4/6-specific inhibitors trilaciclib and palbociclib (Figure 4D). Pre-treatment with both inhibitors suppressed the *Ifnb1* and *Rsad2* super-induction during TNF stimulation, although the effect of trilaciclib was more prominent (Figure 4E and Figure 4-figure supplement 1A). The trilaciclib pretreatment also significantly reduced the ROS production, LPO synthesis and accumulation (Figure 4F). Accordingly, we found that the trilaciclib treatment during TNF stimulation enhanced the Fth1 and Ftl1 protein expression (Figure 4G) and prevented the intracellular labile iron pool (LIP) increase (Figure 4H). The Hif1a protein levels also increased during the trilaciclib plus TNF treatment (Figure 4I). Perhaps, the latter effect could be mediated via the LIP reduction, because this effect could be reproduced by iron chelator DFP (Figure 3P), and the CDK4/6 inhibitor alone induced neither the Hif1a mRNA nor the protein Hif1a levels (Figure 4I and Figure 4-figure supplement 1B). GPX4 expression remained unaffected by trilaciclib pretreatment (Figure 4-figure supplement 1C). In contrast to IFN*γ*, the trilaciclib pretreatment upregulated the expression of lipid biosynthesis genes *Srebf2*, *Scd2* and *Dhcr24* either alone or during TNF stimulation (Figure 4J). Thus, CDK4/6 inhibition using trilaciclib partially recapitulated the effects of IFN*γ* priming by regulating iron metabolism and, thus, prevented the development of the pathological activation state (PAS) in *sst1*-susceptible macrophages during TNF stimulation.

To investigate the effects of RAR activation on PAS, we pre-treated B6.Sst1S BMDMs prior to TNF stimulation with a retinoic acid derivative, All-Trans Retinoic Acid (ATRA) (Figure 4K). The ATRA pretreatment did not decrease the ROS levels, but completely prevented the 4-HNE adducts accumulation (Figure 4L), and partially reduced the *Ifnb1* and *Rsad2* mRNA levels (Figures 4M). In contrast to trilaciclib, however, the effects of ATRA on LIP and ferritin subunits expression were minimal (Figures 4N and 4O, respectively). Searching for the mechanisms of RAR-mediated LPO inhibition, we found that ATRA pretreatment upregulated a major lipid peroxidation inhibiting enzyme GPX4 (Figure 4P), whose defective induction by TNF in the *sst1*-susceptible macrophages we previously reported (Shivraj M. Yabaji et al., 2025). In addition, ATRA treatment alone or in combination with TNF significantly increased the expression of an anti-ferroptotic gene *Scd2* that promotes the accumulation of the lipid peroxidation resistant monounsaturated fatty acids (MUFA) in cell membranes and a cholesterol biosynthesis gene *Dhcr24* (Figure 4Q). These effects demonstrate that RAR activation remodel macrophage lipid metabolism to increase their resilience to lipid peroxidation-induced damage.

Taken together, our data demonstrate that the cSTAR-predicted IFN*γ* mimetics partially recapitulate effects of IFN*γ* via diverse mechanisms, whose activities converge on protecting activated macrophages from the damage inflicted by lipid peroxidation.

### 4. CDK4/6 Inhibition and RAR Activation Cooperate to Prevent pPAS and Improve Intracellular Mtb Control at Lower Doses

To understand how CDK4/6- and RAR-mediated pathways interact during in macrophage activation, we analyzed the structure of the cSTAR-inferred network (Rukhlenko et al., 2022). cSTAR leverages Modular Response Analysis (MRA) (Kholodenko et al., 2021) for both network inference and predicting network function. First, we used cSTAR to infer the network that regulates macrophage activation and demonstrated the overall positive effect of RAR and negative effect of CDK4/6 (Figure 5A). To understand the operation of the inferred network we analyzed the cSTAR-predicted global response matrix (Figure 5B). The global response matrix predicts the change in node activity following a 1% activation of a target node (Kholodenko et al., 2021). Hierarchical clustering was applied to identify co-activated or mutually antagonistic nodes (Figure 5B). Here, columns show the responses of the entire network to a specific node’s activation, whereas rows show a single node’s response to the activation of all other nodes. Hierarchical clustering of the response matrix reveals two mutually antagonistic clusters. The first, comprising RAR, mTOR, and IFN*γ* signaling modules, shows mutual intra-cluster activation; notably, activating these nodes increases Mtb resistance. In contrast, the second cluster (PAK, CDK1/2, and CDK4/6) promoted Mtb susceptibility, also in agreement with DPD_TB_ score predictions (Figure 4C). Therefore, we focused on the interplay between CDK4/6, the strongest contributor to Mtb susceptible phenotype, and key contributors to Mtb resistant phenotype - IFN*γ* and RAR, that clustered closest to IFN*γ* in the network.

**Figure 5:**
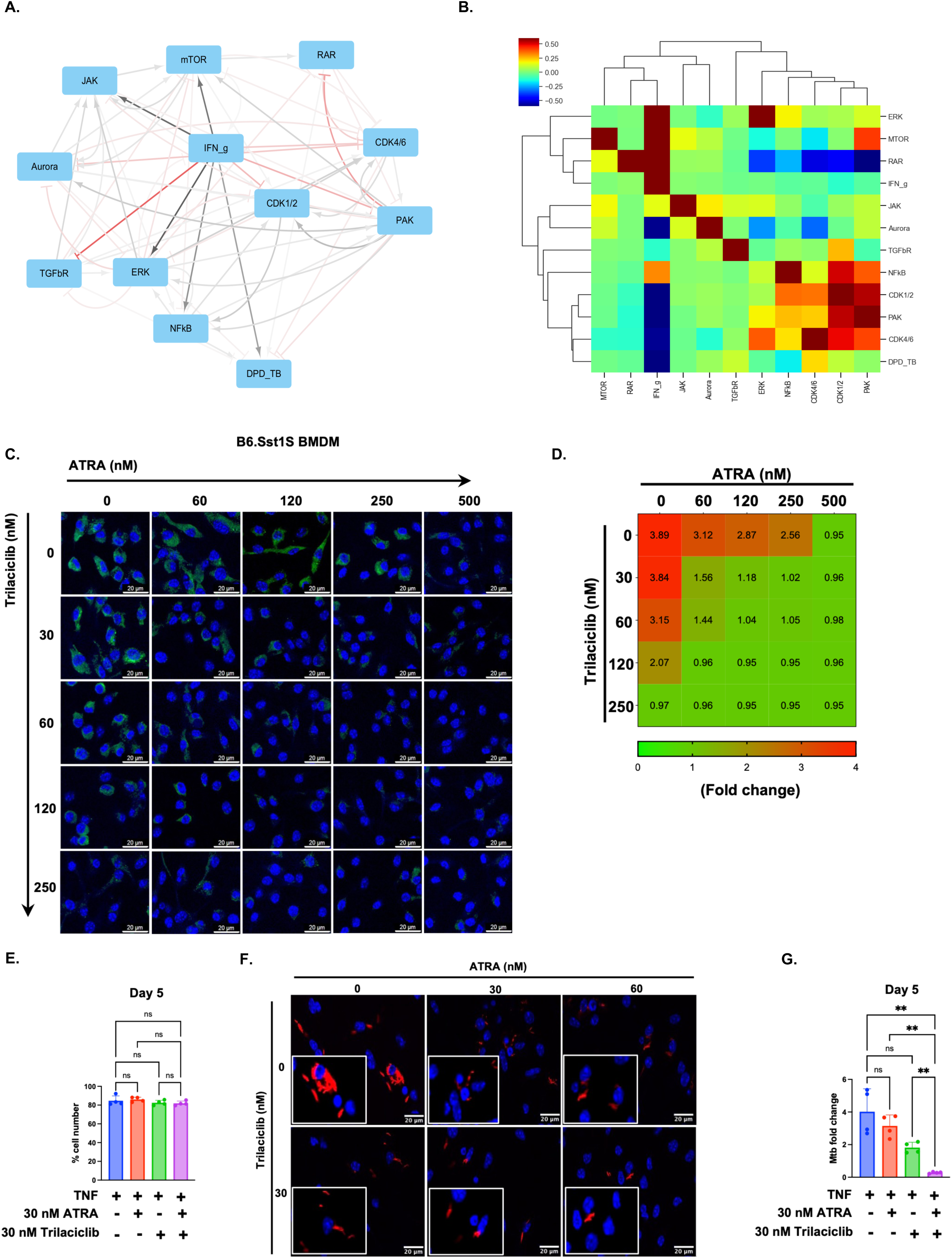
The combination of CDK4/6 inhibition and RAR activation allows dose reduction while preventing pPAS and controlling intracellular Mtb growth. A. cSTAR-inferred network that controls resistance priming of macrophages. Black arrows indicate activation, while red arrows indicate inhibition. The intensity of the color is proportional to the strength of connection. Perturbation data for THP1 cell line were used for network inference. B. Hierarchical clustering of the cSTAR-predicted global response matrix. Each column of the heatmap shows responses of all nodes to activation of the node in question, while each row of the heatmap shows responses of the node in question to activation of all other nodes. Correlation distance was used for the clustering algorithm. C. The combination of trilaciclib and ATRA enables dose reduction effectively suppressing 4-HNE adducts accumulation during TNF stimulation. B6.Sst1S BMDMs were pretreated with Trilaciclib and ATRA at different concentration combinations for 12 h followed by TNF (10 ng/mL) for 36 h. Accumulation of 4-HNE adducts was evaluated by confocal microscopy using a 4-HNE-specific antibody. Scale bar-20 *μ*m. D. Heat map showing the average fold change in 4-HNE adducts fluorescence upon Trilaciclib and ATRA combinations compared to the untreated samples. (n=3). E-G. The combination of 30 nM of trilaciclib and 30 nM ATRA did not cause B6.Sst1S BMDMs cell death during Mtb infection. (n=4). The combination of trilaciclib and ATRA controls the intracellular Mtb growth. B6.Sst1S BMDMs were pretreated with trilaciclib (30 nM) and ATRA (30 nM or 60 nM) alone or in combination for 12 h followed by TNF (10 ng/mL) for 24 h. Following day, cells were infected with Mtb Erdman reporter strain (SSB-GFP, smyc’::mCherry) at MOI=1 for five days. Day 1 and 5 post infection, total cell numbers (E) were quantified by automated microscopy and intracellular Mtb load was quantified by qPCR based assay (G). Images of infected BMDMs were acquired by confocal microscopy 5 days post infection (F). Scale bar: 20 μm. (n=4). The data are presented as mean ± standard deviation (SD) from three-five samples per experiment, representative of three independent experiments. The statistical analysis was performed by one-way ANOVA with Tukey’s multiple comparisons test (Panel E and G). Significant differences are indicated with asterisks (ns, non significant; *, p < 0.05; **, p < 0.01; ***, p < 0.001; ****, p < 0.0001).

We analyzed how activation of each of these 3 nodes affects activity of the other 2 key nodes. It can be seen in Figure 5B, the activation of CDK4/6 suppresses activity of both RAR and IFN*γ*, and activation of IFN*γ* and RAR suppresses activity of CDK4/6. This behavior is caused by mutually inhibiting connections between CDK4/6 and RAR, and between CDK4/6 and IFN*γ* (Figure 5A). Such network topology is known to generate bi-stable states, with either one or the other node to be active, but not both simultaneously (Tsyganov et al., 2012). Based on the network topology, we hypothesized that combination of CDK4/6 inhibition and RAR stimulation could produce synergistic effect in inducing the Mtb resistant macrophage activation state.

To experimentally test the predicted synergy of CDK4/6 inhibition and RAR activation on pPAS, we pretreated B6.Sst1S macrophages individually with various concentrations of trilaciclib, ATRA and their combinations followed by TNF stimulation and measured the 4-HNE adducts accumulation (Figure 5C, 5D). At higher concentrations, treatments with either trilaciclib (250 nM) or ATRA (500 nM) prevented the 4-HNE adducts accumulation. Their combination, however, was efficient at limiting the TNF-induced lipid peroxidation at almost 10 times lower concentrations of trilaciclib (30 nM) and ATRA (60 nM).

Next, we evaluated whether this combinatorial treatment could improve control of intracellular Mtb growth in B6.Sst1S macrophages. Of note, treatments of Mtb-infected BMDMs with the higher concentrations of trilacilib (1.5 μM) or ATRA (2μM) during the 5 day-long experiment caused macrophage death (Figure 5-figure supplement 1A and 1B). The lower doses of trilaciclib (30 nM) or ATRA (30 or 60 nM), however, were not toxic (Figure 5E, Figure 5-figure supplement 1C) and partially reduced the intracellular Mtb growth (Figure 5F, and Figure 5-figure supplement 1D). Notably, the combination of trilaciclib and ATRA at low concentrations preserved the infected macrophage viability and further reduced the intracellular Mtb loads (Figure 5E, 5G). In fact, in this treatment groups the Mtb loads at Day 5 post infection were lower as compared to Day 1, which is suggestive of Mtb clearance by the susceptible macrophages (Figure 5-figure supplement 1E and 1F). Thus, combining trilaciclib and ATRA at low concentrations to treat susceptible macrophages prevented the pathological activation state, preserved their viability and significantly improved their ability to control Mtb.

## Discussion

Our findings reveal a fundamental divergence between type I and type II interferon programs in macrophages. They suggest that the conflicting roles of the type I and type II interferons during chronic TB can be accounted for by their opposing effects on lipid peroxidation: IFN-I enhances, while IFN*γ* prevents it (Table 1). We found that persistent IFN-I signaling promotes a self-amplifying cycle of ROS production and lipid peroxidation that impairs host resistance to Mtb, whereas IFN*γ* primes macrophages for antimicrobial activation while simultaneously increasing their resilience to oxidative damage through iron sequestration, thereby limiting the conversion of ROS into damaging lipid peroxidation products. Thus, beyond differences in downstream effector pathways, IFN-I and IFN*γ* differ in their ability to couple inflammatory activation to cyto-protection, a distinction with important biological causes and pathological consequences.

**Table 1:** Differential regulation of pPAS features by interferons.

| | | IFNβ<br>(during TNF or alone) | IFNγ |
| --- | --- | --- | --- |
| IFN-I signaling | Ifnb | ↑↑ | = |
|  | Rsad2 | ↑↑ | = |
|  | Acod1 | ↑ | ↑ |
| Iron metabolism | Labile iron pool | = | ↓ |
|  | FTH1 | = | ↑↑ |
|  | FTL | = | ↑↑ |
| Oxidative stress & lipid peroxidation | ROS | ↑ | ↑ |
|  | LA alkyne | ↑ | ↑ |
|  | 4-HNE | ↑ | = |
| Lipid biosynthesis | Srebf2 | ↓ | ↓ |
|  | Scd2 | ↓ | ↓ |
|  | Dhcr24 | ↓ | ↓ |
| Net result | pPAS | ↑ | ↓ |

↑ Induction level
↓ Suppression level
= Not affected

We demonstrate that persistent IFN-I signaling and autocatalytic lipid peroxidation form a self-reinforcing pathological circuit, in which ROS-driven lipid damage further amplifies inflammatory signaling leading to the IFN-I pathway hyperactivity. This circuit is sufficient to maintain the persistent pathological activation state (pPAS) of macrophages in an autocrine manner after removal of TNF, the initial PAS trigger in our in vitro model. The pPAS is characterized by the continuous ROS generation, the accumulation of lipid peroxidation products, suppression of lipid biosynthesis, and further escalation of the IFN-I pathway upon repeated TNF stimulation. Mechanistically, this vicious cycle develops in *sst1* susceptible macrophages because of their failure to upregulate the *sst1*-encoded IFN-inducible proteins SP110 and SP140, which serve as feedback regulators of the IFN-I pathway either directly (Ji et al., 2021; Witt et al., 2025) or indirectly, downstream of Myc hyperactivity (Shivraj M. Yabaji et al., 2025). We propose that other molecular pathologies characterized by persistent cellular stress and IFN-I hyperactivity, such as interferonopathies, may be driven by similar IFN-I/lipid peroxidation amplification loops.

Because type I interferons are produced by virus-infected cells, as well as by cells undergoing chronic stress or damage (Boccuni et al., 2022), we hypothesize that the IFN-I/lipid peroxidation amplification loop may represent an element of an evolutionarily conserved host-defense strategy for containing or eliminating infected, damaged, or otherwise aberrant cells. Consistent with this hypothesis, toxic aldehydes generated by lipid peroxidation have been shown to limit the replication of viruses and intracellular pathogens (Anaya-Sanchez et al., 2025), and IFN-I signaling has been implicated in the elimination of epigenetically altered or premalignant cells in vitro (Leonova et al., 2013; Zhao et al., 2021). Similarly, lipid peroxidation-induced cellular senescence could restrict aberrant cell proliferation. We found that IFN-I promotes lipid peroxidation by inducing ROS without iron sequestration. This mode permits ROS amplification and may be more efficient in containing acute infection. However, during chronic infection or persistent inflammation, sustained activation of this pathway becomes maladaptive. In chronic TB lesions, this response is particularly detrimental because the lipid-rich cell wall and non-replicative persistence programs of Mtb allow the bacterium to withstand the initial host attack, whereas host macrophages are vulnerable to self-inflicted oxidative injury. The resulting macrophage damage impairs their antimicrobial effector functions and antigen presentation thereby creating a permissive niche for the bacterial persistence and growth. This framework provides a plausible explanation for the dual role of type I interferon in TB (Desvignes et al., 2012; Munson & Kaushal, 2025).

In contrast, IFN*γ* is produced not by the infected or stressed cells themselves, but by specialized immune cells, primarily T lymphocytes and NK cells, upon recognition of specific targets. Thus, IFN*γ* functions as a paracrine messenger that alerts and preconditions host tissues before or during pathogen encounter. Its local and systemic pleiotropic effects include priming macrophages for subsequent activation through epigenetic and metabolic reprogramming, enabling them to reallocate cellular resources toward antimicrobial effector functions. Our current studies reveal that, in parallel with boosting ROS production, IFN*γ* enhances macrophage resilience to oxidative stress by limiting the labile iron pool, thereby preventing the amplification of ROS-mediated host-cell damage through Fenton chemistry. By coordinating antimicrobial effector mechanisms with anticipatory cyto-protection, IFN*γ* limits self-inflicted oxidative injury during full-scale activation, supports more robust and sustained effector activities, and reduces bystander tissue damage. This mechanism may help explain the paradoxical anti-inflammatory and tissue-protective effects of this classical pro-inflammatory cytokine in vivo.

Both type I and type II interferon pathways evolved as host defense mechanisms against taxonomically diverse pathogens. Conceivably, their cross-regulation and mutual antagonism help shape responses appropriate for viral versus intracellular bacterial infections, respectively. However, during Mtb infection lesion dynamics and architecture critically influence the local IFN balance. The IFN-I-mediated protection is transient and limited to the early stages of infection, when and where macrophage - Mtb interactions occur primarily in a cell autonomous manner. Following adequate T cell activation, IFN*γ*-mediated macrophage reprogramming provides a more efficient coordinated response, enabling both control of intracellular Mtb and tissue protection. Consistent with this model, BCG vaccination (Yan et al., 2007) or prior Mtb infection (Gern et al., 2025) accelerate the formation of IFN*γ*-dominated protective lesions, even in the genetically susceptible but immunocompetent mice.

However, in the susceptible animals, preexisting immunity delays, but does not prevent lesion progression (Cohen et al., 2022; S. M. Yabaji, S. Lata, A. E. Tseng, et al., 2025; Yan et al., 2007). Myeloid cells increasingly dominate TB lesions, whereas T cells are relegated to the lesion periphery (Branchett et al., 2025; McCaffrey et al., 2026; McCaffrey et al., 2022; S. M. Yabaji, S. Lata, A. E. Tseng, et al., 2025). As expanding myeloid compartments spatially segregate infected macrophages from IFN*γ*-producing lymphocytes, the local interferon balance shifts toward IFN-I dominance, because Mtb-infected and aberrantly activated stressed macrophages produce IFNγ in a cell autonomous manner. This shift leads to gradual loss of protection within individual lesions and promotes hyperinflammatory tissue damage. Eventually, bacteria resume intracellular replication, destroy host cells, and begin growing extracellularly. Subsequent neutrophil recruitment expands local tissue damage, promotes bacterial expansion (Eruslanov et al., 2005), and further stimulates the IFN-I pathway (Kotov et al., 2023). In addition, systemic IFN-I hyperactivity compromises IFN*γ*-mediated hematopoietic reprogramming (Kaufmann et al., 2018; Khan et al., 2020). Thus, in susceptible hosts the immune system becomes locked into an IFN-I-dominant state at both local and systemic levels.

Preventing macrophages from transitioning into an IFN-I-dominant pathological activation state may therefore represent a logical therapeutic strategy. We searched for small molecules that could recapitulate protective components of IFN*γ*-mediated macrophage reprogramming and promote an oxidative-stress-resilient, Mtb-resistant state.

Using cSTAR analysis of resistant and susceptible macrophage states, we found that IFN*γ* priming shifted B6.Sst1S susceptible macrophages toward the transcriptional state of B6 resistant macrophages. We then confirmed that this transcriptome shift is closely correlated with an increased ability of susceptible macrophages to control Mtb replication. Thus, using transcriptomic data, we have built DPD_TB_ score as a novel in vitro metric for macrophages priming to be either susceptible or resistant to Mtb infection, consistent with the in vivo phenotype.

Next, we used publicly available data to identify perturbations that could regulate the transition of susceptible macrophages toward the IFN*γ*-mediated Mtb-resistant state. This analysis identified several candidate pathways predicted to either mimic or antagonize the effects of IFN*γ*. However, the pathways that increased macrophage resistance only partially recapitulated the potent effect of IFN*γ* priming. Therefore, we assessed interactions among the candidate pathways and identified two mutually antagonistic regulatory clusters. The Mtb resistance-promoting cluster comprised retinoic acid receptor (RAR), mTOR, and IFN*γ* signaling modules, which showed mutual intra-cluster activation. The antagonistic cluster, consisting of PAK, CDK1/2, and CDK4/6 nodes, was predicted to promote Mtb susceptibility. We focused on the interplay between CDK4/6, the strongest contributor to the Mtb-susceptible phenotype, and RAR, a key contributor to the Mtb-resistant phenotype. Network topology analysis suggested that simultaneous CDK4/6 inhibition and RAR stimulation could synergize to induce an Mtb-resistant macrophage activation state.

The importance of retinoic acid (RA) in host resistance to tuberculosis has been demonstrated in humans and animal models (Faye Lanni, 2025; Podell et al., 2022). However, the specific mechanisms underlying RA-mediated protection in TB remain incompletely understood. RA has previously been shown to induce expression of interferon regulatory factor 1 (IRF1), a master transcriptional regulator downstream of IFN*γ* receptor signaling, through a STAT1-independent mechanism (Luo & Ross, 2006). In addition, cooperation between retinoic acid receptor (RAR) and IRF1 transcription factors has been reported to enhance transcriptional activation at selected promoters (Clarke et al., 2004). Our studies suggest that, in addition to boosting macrophage effector functions, RA signaling increases macrophage resilience to stress, thereby partially mimicking the ability of IFN*γ* to prevent runaway lipid peroxidation. Unlike IFN*γ*, however, this protective effect did not depend on upregulation of ferritins and was more likely associated with the induction of the anti-ferroptotic enzyme GPX4 and increased lipid biosynthesis.

CDK4/6 inhibition emerged as a second, mechanistically distinct route to an IFN*γ*-like protective state. The role of cyclin-dependent kinases in TB pathogenesis remains poorly understood. However, prior studies using broad-spectrum tyrosine kinase inhibitors, such as imatinib and dasatinib, demonstrated improved outcomes of Mtb infection in mice, primarily by reducing inflammatory tissue damage (Napier et al., 2011; Shee et al., 2026). CDK4/6 are central regulators of cell-cycle progression that are activated by D cyclins in response to mitogenic signals. They phosphorylate Rb and thereby promote downstream regulators of G0-to-S phase progression, including E2F and Myc. CDK4/6 activity also antagonizes terminal differentiation of myeloid progenitors, and CDK4/6 inhibitors have been explored therapeutically in myeloid leukemias (He et al., 2017; Xie et al., 2023).

Our previous work demonstrated increased activity of E2F and Myc pathways in TNF-stimulated B6.Sst1S macrophages compared with B6 wild-type macrophages. Moreover, hyperactivity of the Myc pathway in blood transcriptomes from TB patients was associated with poorer treatment outcomes (Shivraj M. Yabaji et al., 2025). Of note, recent in vivo single-cell RNA-seq studies in genetically resistant and susceptible mice after aerosol Mtb infection demonstrated that increased activity of Myc and E2F pathways was among the earliest manifestations of *sst1*/Sp140-mediated susceptibility, preceding IFN-I pathway upregulation (Duffy et al., 2026). We found that CDK4/6 inhibitors recapitulated the ability of IFN*γ* to stimulate ferritin expression and reduce the intracellular labile iron pool in an *sst1*-independent manner. This reallocation of intracellular iron into ferritin storage pools likely reflects a transition away from an anabolic, proliferation-supporting state toward terminal differentiation and priming for activation. As discussed above, conversion of labile redox-active iron into redox-inactive ferritin stores would limit ROS-driven Fenton chemistry and thereby reduce oxidative host-cell damage.

Together, our findings suggest a strategy to reduce inflammatory pathology without broadly suppressing host defense. They support a model in which the outcome of interferon signaling in TB depends on whether inflammatory activation is coupled to oxidative-stress resilience. In *sst1*-susceptible macrophages, persistent autocrine IFN-I signaling promotes ROS production, lipid peroxidation, suppression of lipid repair, and further IFN-I amplification, producing a self-reinforcing pathological activation state. In contrast, IFN*γ* acts as a paracrine anticipatory signal that primes macrophages for antimicrobial function while limiting labile iron and lipid peroxidation, thereby preserving cell viability and effector capacity. Pharmacological activation of RAR signaling and inhibition of CDK4/6 recapitulate complementary components of this IFN*γ*-protective state (Table 2) and may jointly better resist pathological activation state. Importantly, combining these small molecules may allow lower doses of each agent, potentially enhancing efficacy while mitigating nonspecific toxicity. We anticipate that such disease-modifying approaches, used together with antibiotics, may reduce inflammatory tissue damage, shorten treatment duration, and limit post-TB lung impairment and systemic inflammatory complications. More broadly, these findings suggest that pharmacological reinforcement of oxidative-stress resilience may represent a strategy for limiting interferon-associated tissue pathology in chronic inflammatory disease

**Table 2:** IFN*γ* mimicking effect of trilaciclib and ATRA on pPAS features during TNF stimulation.

|  |  | **IFNγ** | **Trilaciclib** | **ATRA** |
| --- | --- | --- | --- | --- |
| **Oxidative stress** | **ROS** | ↑ | ↑ | ↑ |
| **Iron metabolism** | **Labile iron pool** | ↓↓ | ↓↓ | ↓ |
|  | **FTH1** | ↑↑ | ↑↑ | ↑ |
|  | **FTL** | ↑↑ | ↑↑ | ↑ |
| **Lipid peroxidation** | **GPX4** | = | = | ↑ |
|  | **LA alkyne** | ↓ | ↓ | NT |
|  | **4-HNE** | ↓ | ↓ | ↓ |
| **Lipid biosynthesis** | **Srebf2** | ↓ | ↑ | ↑↑ |
|  | **Scd2** | ↓ | ↑ | ↑↑ |
|  | **Dhcr24** | ↓ | ↑ | ↑↑ |
| **IFN-I signaling** | **Ifnb** | ↓ | ↓ | ↓ |
|  | **Rsad2** | ↓ | ↓ | ↓ |
| **Net result** | **pPAS** | ↓ | ↓ | ↓ |

↑ Induction level
↓ Suppression level
= Not affected

## Methods

### Cell culture

Bone marrow cells were isolated from the tibia and fibula of either B6 or B6.Sst1S mice. The bone marrow cell isolation, culture, and bone marrow-derived macrophage (BMDM) generation were performed according to the method previously described (Yabaji et al., 2022).

### Determination of lipid peroxidation product synthesis/Click-iT lipid peroxidation detection

BMDMs were cultured on cover slips in 24-well plates. After the treatments, linoleamide alkyne (LAA) was added to the culture at a concentration of 50 μM 1 h prior to cell harvest. Cells were processed according to the instructions provided in the manual. Briefly, cells were washed with PBS and fixed with 3.7% formaldehyde in PBS for 15 min at room temperature. Cells were washed with PBS and permeabilized with 0.5% Triton X-100 in PBS for 10 min at room temperature. Further, blocking was carried out with 1% BSA in PBS for 30 min at room temperature. Cells were washed with PBS and incubated with Click-iT reaction cocktail for 30 min at room temperature, protected from light. Cells were washed with 1% BSA in PBS and further with PBS. The coverslip was mounted onto the slide using Prolong Gold antifade reagent with DAPI. The fluorescent images were captured using a Leica SP5 confocal microscope. The images were analyzed using ImageJ software (NIH).

### Immunofluorescence staining for 4-HNE adducts

BMDMs were cultured on cover slips in 24-well plates. After the treatments, fixation, permeabilization, and blocking were performed as mentioned in the earlier sections. The cells were incubated with the 4-HNE antibody overnight at 4°C. The cells were washed with 1% BSA in PBS and incubated with Alexa Fluor 488 conjugated goat anti-rabbit antibodies for 1 h at room temperature, protected from light. The cells were washed with PBS, and the coverslip was mounted onto the slide using Prolong Gold antifade reagent with DAPI. The fluorescent images were captured using a Leica SP5 confocal microscope. The images were analyzed using ImageJ software (NIH).

### Intracellular measurement of labile iron pool

BMDMs were cultured in 96-well plates. After the treatments, cells were washed with serum-free culture medium and incubated with 1 mM FerroOrange solution in serum-free media for 30 min at 37°C and 5% CO₂. The fluorescent images were captured using the Perkin-Elmer Operetta CLS High-Content Analysis System, and the signal quantification was performed using software.

### Measurement of reactive oxygen species

BMDMs were cultured in 96-well plates. After the treatments, CellROX Green reagent was added to the cell culture at a concentration of 5 mM and further incubated for 30 min at 37°C and 5% CO₂. The cells were washed with PBS and fixed using 3.7% formaldehyde in PBS for 15 min at room temperature. The cells were washed with PBS and resuspended in PBS. The fluorescent images were captured using a Nexcelom Celigo Micro-Well Plate Imager and analyzed using ImageJ software (NIH).

### Real-time quantitative RT-PCR

The BMDMs were cultured in 6-well plates. After the treatments, cells were washed with PBS and collected in TRIzol reagent. The total RNA was isolated using the RNeasy Plus Mini Kit (QIAGEN), involving the DNase treatment using RNase-Free DNase (QIAGEN) following the recommended procedure. The total RNA concentration was quantified using the NanoDrop 1000 Spectrophotometer. The cDNA synthesis was carried out from 500 ng of total RNA from each sample using PrimeScript RT Master Mix (Takara Bio USA) following the recommended protocol. The qRT-PCR was performed using the reagents of GoTaq qPCR Master Mix (Promega Corporation) and the CFX Opus 96 Real-Time PCR system (Bio-Rad Laboratories). The thermal cycle conditions are as follows: 95°C for 3 min followed by 39 cycles of 95°C for 15 sec and 55°C for 1 min and finally the melt curve conditions as preset in the machine. The fold change in gene expression was calculated by the ddCT method using either 18S rRNA or β-actin as endogenous control.

### RNA-seq library preparation & sequencing

B6 and B6.Sst1S BMDMs were treated with TNF, IFN*γ*, and in combination of both for 12 h and total RNA was extracted using TRIzol reagent as mentioned in previous section. Total RNA integrity was verified using the High Sensitivity RNA Pico 6000 assay run on an Agilent 2100 Bioanalyzer (Agilent Technologies, Palo Alto, CA). To enrich for mRNA, 150 ng of total RNA for each sample was incubated with magnetic Oligo-d(T)_25_ beads per the manufacturer’s instructions using NEB Next Poly(A) mRNA Magnetic Isolation Module (New England Biolabs, USA). Libraries were prepared using NEB Next Ultra II Directional RNA Library Kit for Illumina according to the manufacturer’s protocol. Briefly, after mRNA enrichment, the RNA was fragmented at 94°C for 15 min. First strand cDNA was generated, and incorporation of dUTP occurred during synthesis of second strand cDNA to indicate strandedness. Double-stranded cDNA then underwent end repair, A-tailing, adaptor ligation, and incorporation of sample-specific multiplex indices via PCR amplification (11 cycles) according to the manufacturer’s protocol (New England Biolabs, USA). Size distribution and molarity of amplified cDNA libraries were assessed via the Bioanalyzer High Sensitivity DNA Assay (Agilent Technologies, USA). All cDNA libraries were sequenced on an Illumina NextSeq 2000 instrument at 600 pM with 2% PhiX control library spiked in (Illumina, USA) using P3 100 cycles sequencing kit (Illumina, USA) and resulting in 55-91 million reads per sample (50x50bp paired-end reads). All samples were performed in triplicates.

### RNA-seq pre-processing and differential expression analysis

Data pre-processing and the downstream computational analyses were all performed using R statistical software v4.4.3 (Core, 2022). Genes with lower total counts of 10 across samples were removed. RNA counts were normalized with DESeq2 and log-2-transformed (Love et al., 2014). The data included two biological factors: genotype (wild type and B6.Sst1S) and treatment (untreated, TNFα, IFN*γ*, and TNFα+IFN*γ*). To identify differentially expressed genes (DEGs), contrasts were defined as follows:

1. Within-genotype treatment contrasts: comparisons between all possible treatments within each genotype (e.g., TNFá+IFN*γ* vs TNFα in B6.Sst1S or wild type).
2. Treatment contrasts accounting for genotype: comparisons between all possible treatments with genotype as a covariate.
3. Genotype contrasts accounting for treatment: comparisons between all possible genotypes with treatment as a covariate.

Differentially expressed genes were filtered using a standard threshold (absolute fold-change > 1.5 and BH adjusted p-value < 0.05).

### Principal component analysis

Principal component analysis was performed for all samples using the PCAtools R package v2.16 (Blighe K, 2025). Colour represents treatment type and shape represents genotype.

### Gene set enrichment and semantic reduction analysis

Differential expression analysis statistic scores were used for gene set enrichment analysis using the fgsea R package v1.30 (Korotkevich et al., 2021). The default fgsea parameters were used, except for the minSize parameter, which was set to 15. The gene ontology biological processes (GO:BP) database, obtained from the Human Molecular Signature Database (MSigDB), was used for pathways reference (Ashburner et al., 2000; Liberzon et al., 2015). Significant enriched pathways were defined with adjusted p-value < 0.05. Semantic reduction of significantly enriched pathways were performed with the rrvgo R package v1.16 (Sayols, 2023).

### Single sample gene set enrichment analysis

Single sample gene set enrichment analysis were performed with the GSVA R package v1.52 (Hanzelmann et al., 2013). Several pathways were picked to be further explored that were obtained either from GO:BP or Hallmark (Ashburner et al., 2000; Liberzon et al., 2015). Specifically, for ferroptosis-induction and ferroptosis-inhibition pathways, the underlying genes were obtained from Amaral et.al study (Amaral et al., 2024). The gene list for each pathway was included in Data file 1.

### cSTAR analysis

To build State Transition Vector STV_TB_ we have applied ordinary least squares linear regression of transcriptomic data for cytokine stimulation (Figure 4A) over Mtb fold change data (Figure 3A). Since the available perturbation transcriptomics data from CMAP dataset had only 978 genes measured directly (Subramanian et al., 2017), we have built STV_TB_ in the space of these 978 genes. To reflect increase of TB resistance in response to the increase of appropriate DPD score, DPD_TB_, we have multiplied STV_TB_ on -1, so the increase in DPD_TB_ corresponds to the decrease in Mtb fold change, and consequently, increase in TB-resistance.

Next, we utilized publicly available perturbation data to understand how priming of macrophages towards Mtb resistance can be controlled. Particularly, we used a cSTAR pipeline adapted for perturbation transcriptomics data, that allows reconstruction of the protein signaling network wiring from transcriptomics responses perturbation data (Deng et al., 2025). This approach allows us to access how pharmacological interventions targeting different druggable proteins move macrophage states along the STV_TB_, by changing DPD_TB_ score. Among cellular models that are considered relevant for the task of controlling macrophage cell states, THP1 cell line has the highest amount of perturbation data. Accordingly, we have collected perturbation data for this cell line to run our cSTAR pipeline (Deng et al., 2025). While most of perturbation data that we used originated from CMAP dataset (Subramanian et al., 2017), this dataset lacks perturbations to some critically important nodes, such as RAR receptors and receptors of IFN*γ*. For these nodes we have taken data from the following GEO accession numbers: GSE46268, GSE244129 and GSE229710. Since THP1 cells do not directly resemble *sst1*-driven susceptibility phenotype, we excluded IFN type I and TNF from the network, and focused on other key signaling nodes.

In brief, we collated from different publicly available sources (see above) a dataset of perturbation responses, combining log fold changes of gene expression levels with respect to appropriate control. For each data point, we have calculated DPD_TB_ responses to specific perturbation. Next, we have selected core network components based on ranking of DPD_TB_ responses, as well as on our prior knowledge of key pathways involved. Then, we inferred module activity coefficients, which allow calculating module activities, as described in (Deng et al., 2025). Using calculated module activities, we have calculated matrix of global response coefficients *R*, that served as an input to Bayesian Modular Response Analysis (BMRA) (Tapesh Santra, 2018) method, and inferred matrix of local connections *r*. According to MRA theory, each column of −*r*^-1^ matrix represents MRA-predicted responses of all network nodes to 1% activation of the specific node (Kholodenko et al., 2021). Thus, we plotted elements of −*r*^-1^ matrix as cSTAR network predictions at Figures 4C and 5B.

### Bacterial culture

Mtb Erdman reporter strain (SSB-GFP, smyc::mCherry) was grown in Middlebrook 7H9 broth supplemented with 10% Oleic Albumin Dextrose Catalase (OADC) in a shaking incubator at 37°C. For infection experiment, bacteria were grown until the log phase (OD 0.4-0.6), centrifuged at 3000 rpm for 10 min and resuspended in DMEM F/12 supplemented with 10% FBS, 20 mM HEPES, 2 mM L-glutamine, 10% L929 cell conditioned media (LCCM). The bacteria were briefly sonicated using a sonicating water bath at 20°C-25°C. Single bacterial suspension was obtained by passing the bacteria through 5 μm filter using control syringe. The bacterial count was measured at OD_600_ and considered OD_600_ 1 = 3x10^8^ bacteria per ml.

### M. tuberculosis infection of BMDMs

B6 or B6.Sst1S BMDMs were cultured in 96-well plates supplemented with DMEM F/12, 10% FBS, 20 mM HEPES, 2 mM L-glutamine, 10% LCCM. The single bacterial suspension was diluted in the above media without antibiotics to achieve the multiplicity of infection (MOI) - 1. Hundred microliters of the Mtb were added to the BMDMs pretreated with TNF, IFN*γ*, TNF + IFN*γ*, TNF + trilaciclib, or TNF + ATRA. The culture plates were centrifuged at 500 g for 5 min and incubated at 37°C for 1 h. The extracellular bacteria were eliminated by Amikacin treatment at 200 μg/ml for 45 min at 37°C. The cells were washed and cultured in presence of the treatment conditions. Post day 1 and day 5 of infection, cells were harvested and the cell numbers and Mtb load were quantified as described earlier (Yabaji et al., 2022). All experimental procedures involving Mtb were executed within Biosafety Level 3 (BSL-3) containment facilities in strict compliance with Boston University’s Institutional Biosafety Committee (IBC) approved protocol #25-875, as well as biosafety standards established by the Centers for Disease Control and Prevention and the National Emerging Infectious Disease Laboratories.

### Western blotting

BMDMs were washed with PBS and were collected by scraping in the ice-cold lysis buffer (20 mM Tris-HCl. pH 7.5, 150 mM NaCl, 1 mM EDTA, 1.5 mM MgCl₂, 1% NP-40 alternative, 10% glycerol, 50 mM sodium fluoride, 0.1 mM sodium orthovanadate, 1 mM PMSF, 1X protease inhibitor cocktail, 1X phosphatase inhibitor II, 1X phosphatase inhibitor III). The cell lysis was performed by brief sonication, and the cell lysate was collected by centrifugation at 12,000 g for 10 min at 4°C. Total protein concentration was determined by using Bradford reagent (Bio-Rad Laboratories). The Laemmli buffer (2X) was added to each sample, and the proteins were denatured at 95°C for 5 min. The cell lysates were resolved by SDS-PAGE, and the protein bands were electro-blotted onto the PVDF membranes. The membranes were blocked with 3% BSA in TBST and probed with primary antibodies overnight at 4°C. After washing the membranes with TBST, they were further incubated with relevant secondary antibodies. The membranes were washed with TBST, and the chemiluminescent signals were captured using SuperSignal West Pico Chemiluminescent Substrate (Thermo Scientific) in the FujiFilm LAS-4000 Luminescent Image Analyzer. The protein band densities were quantified using the ImageJ software (Version 1.54P). All the experiments were carried out in two biological replicates and beta-tubulin was considered as an endogenous loading control, and the normalized relative band intensities were calculated.

### Statistical analysis

All graphs were plotted, and statistical analyses were performed using GraphPad Prism Version 10.6.1. For comparison across multiple groups based on single variable, ordinary one-way ANOVA with Tukey’s multiple comparisons test, with a single pooled variance was applied. Comparisons among groups involving two or more variables were analyzed by two-way ANOVA with Tukey’s multiple or Sidak tests. Statistical significance levels are indicated as: ns, non-significant; *,p<0.05; **,p<0.01; ***,p<0.001; and ***,p<0.0001.

## Supporting information

Key Resources Table

## Data availability

The code for RNA-seq pre-processing and differential expression analysis is available on GitHub at https://github.com/ZainulArifin1/Boston_Macrophae_Transcriptomics_Analysis/tree/main. The code for cSTAR analysis is available on GitHub at https://github.com/OleksiiR/THP1_TB_cSTAR/. The RNAseq data are available under GEO accession number GSE337186.

## Acknowledgements

This work was supported by NIH grants R01HL126066 to IK, R01AI189410 to WRB and IK, EU grant no. 101136926 MULTIR to BNK, Research Ireland grant 22/PATH-S/10875 to OSR, and Leducq Foundation for Cardiovascular Research grant ARTIST to OSR and BNK. We would like to thank Dr. Michael Kirber, the Boston University Cellular Imaging, and Sequencing Resource cores for their advice and technical assistance. We thank Dr. Lingyi Lynn Deng, Mr. Matthew Au and the Analytical Instrumentation core for their services. OSR and BNK would like to thank Ciardha Carmody and Prof. Jonathan Bond from Systems Biology Ireland UCD for help in processing THP1 perturbation data from CMAP (LINCS L1000) dataset.

## Authors contributions

Conceptualization IK, SMY, PBA, BNK, OSR, WRB

Data curation PBA, OSR, BNK, IK, VZ, MZA, AAG

Formal analysis PBA, MZA, VZ, OSR

Validation PBA, SMY, SL

Investigation PBA, SL, SMY, OSR, MZA

Methodology PBA, SMY, OSR, VZ, YOA

Writing-original draft PBA, IK

Writing-review and editing IK, AAG, BNK, OSR, WRB

Resources, Supervision IK, BNK

Funding acquisition IK, WRB, BNK, OSR, VZ

Project administration IK

---

**Figure 1–figure supplement 1:**
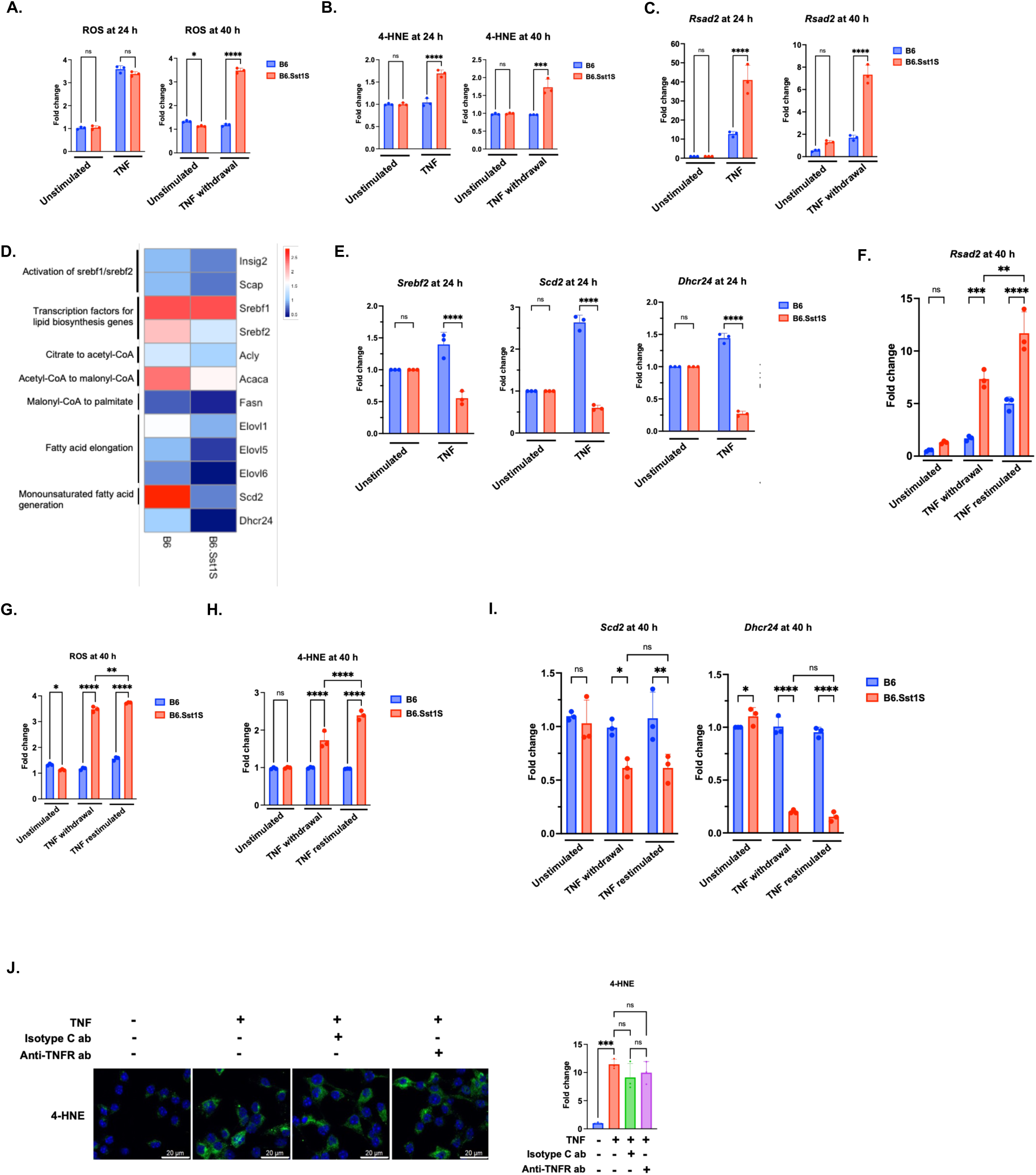
A. ROS levels remain elevated in B6.Sst1S BMDMs following TNF withdrawal. BMDMs derived from B6 and B6.Sst1S mice were treated as described in Figure 1A. Accumulation of ROS was detected using CellROX Green staining, and fluorescence images were acquired by fluorescence microscopy. Bar graphs show the fold change in fluorescence intensity relative to unstimulated B6 controls. B. Accumulation of 4-HNE adducts persists in B6.Sst1S BMDMs following TNF withdrawal. BMDMs from B6 and B6.Sst1S mice were treated as described in Figure 1A. The accumulation of lipid peroxidation product, 4-HNE was detected by confocal microscopy using 4-HNE specific antibody. The 4-HNE accumulation was quantified using ImageJ and plotted as fold accumulation compared to untreated B6 group. C. IFN-I signaling remains persistently super-induced in B6.Sst1S BMDMs following TNF withdrawal. BMDMs from B6 and B6.Sst1S mice were treated as described in Figure 1A. The mRNA expression levels of *Rsad2* were quantified by qRT-PCR at 24 and 40 h post TNF stimulation. Fold change was calculated relative to the unstimulated B6 control using the ΔΔCt method with 18S as an internal control. D. Lipid biosynthesis genes are downregulated in B6.Sst1S BMDMs during TNF stimulation. Heatmap showing the average fold change in FPKM values from RNA-seq analysis of B6 and B6.Sst1S BMDMs following 12 h TNF stimulation. Fold change was calculated relative to unstimulated controls. E. Lipid biosynthesis genes remain downregulated in B6.Sst1S BMDMs following TNF stimulation. BMDMs from B6 and B6.Sst1S mice were treated with 10 ng/mL TNF for 24 h. The mRNA expression levels of *Srebf2, Scd2,* and *Dhcr24* were quantified by qRT-PCR. Fold change was calculated relative to the unstimulated B6 control using the ΔΔCt method with 18S rRNA as an internal control. F. TNF restimulation further enhances IFN-I signaling in B6.Sst1S BMDMs. BMDMs from B6 and B6.Sst1S mice were treated as described in Figure 1A, including TNF restimulation at 24 h. Cells were harvested at 40 h, and *Rsad2* mRNA expression levels were quantified by qRT-PCR. Fold change was calculated relative to the unstimulated B6 control using the ΔΔCt method with 18S as an internal control. G. TNF restimulation during pPAS sustains elevated ROS levels in B6.Sst1S BMDMs. BMDMs from B6 and B6.Sst1S mice were treated as described in Figure 1A, including TNF restimulation at 24 h for additional 16 h. Accumulation of ROS was detected using CellROX Green staining, and fluorescence images were acquired by fluorescence microscopy. Bar graphs show the fold change in fluorescence intensity relative to unstimulated B6 controls. H. TNF restimulation during pPAS further increases 4-HNE adduct accumulation in B6.Sst1S BMDMs. BMDMs derived from B6 and B6.Sst1S mice were treated as described in Figure 1A, including TNF restimulation at 24 h for additional 16 h. The accumulation of lipid peroxidation product, 4-HNE was detected by confocal microscopy using 4-HNE specific antibody. The 4-HNE accumulation was quantified using ImageJ and plotted as fold accumulation compared to untreated B6 group. I. Lipid biosynthesis genes remain suppressed in B6.Sst1S BMDMs during TNF restimulation. BMDMs derived from B6 and B6.Sst1S mice were treated as described in Figure 1A, including TNF restimulation at 24 h for additional 16 h. Cells were harvested at 40 h, and the mRNA expression levels of *Scd2* and *Dhcr24* were quantified by qRT-PCR. Fold change was calculated relative to the unstimulated B6 control using the ΔΔCt method with 18S as an internal control. J. Sustained lipid peroxidation in B6.Sst1S BMDMs is independent of continued TNF signaling. B6.Sst1S BMDMs were stimulated with 10 ng/mL TNF for 24 h. Following the 24 h TNF stimulation, cells were treated with either isotype control antibodies or anti-TNFR antibodies and harvested at 36 h. The accumulation of lipid peroxidation product, 4-HNE was detected by confocal microscopy using 4-HNE specific antibody. The 4-HNE accumulation was quantified using ImageJ and plotted as fold accumulation compared to untreated group. Scale bar: 20 μm. The data are presented as mean ± SD from three-five samples per experiment, representative of three independent experiments. The statistical analysis was performed by two-way ANOVA with Tukey’s multiple comparisons test (Panel A, B, C, and E), Sidak’s multiple comparison test (Panel F, G, H, and I), and one-way ANOVA with Tukey’s multiple comparisons test (Panel J). Significant differences are indicated with asterisks (ns, non significant; *, p < 0.05; **, p < 0.01; ***, p < 0.001; ****, p < 0.0001).

**Figure 2–figure supplement 1:**
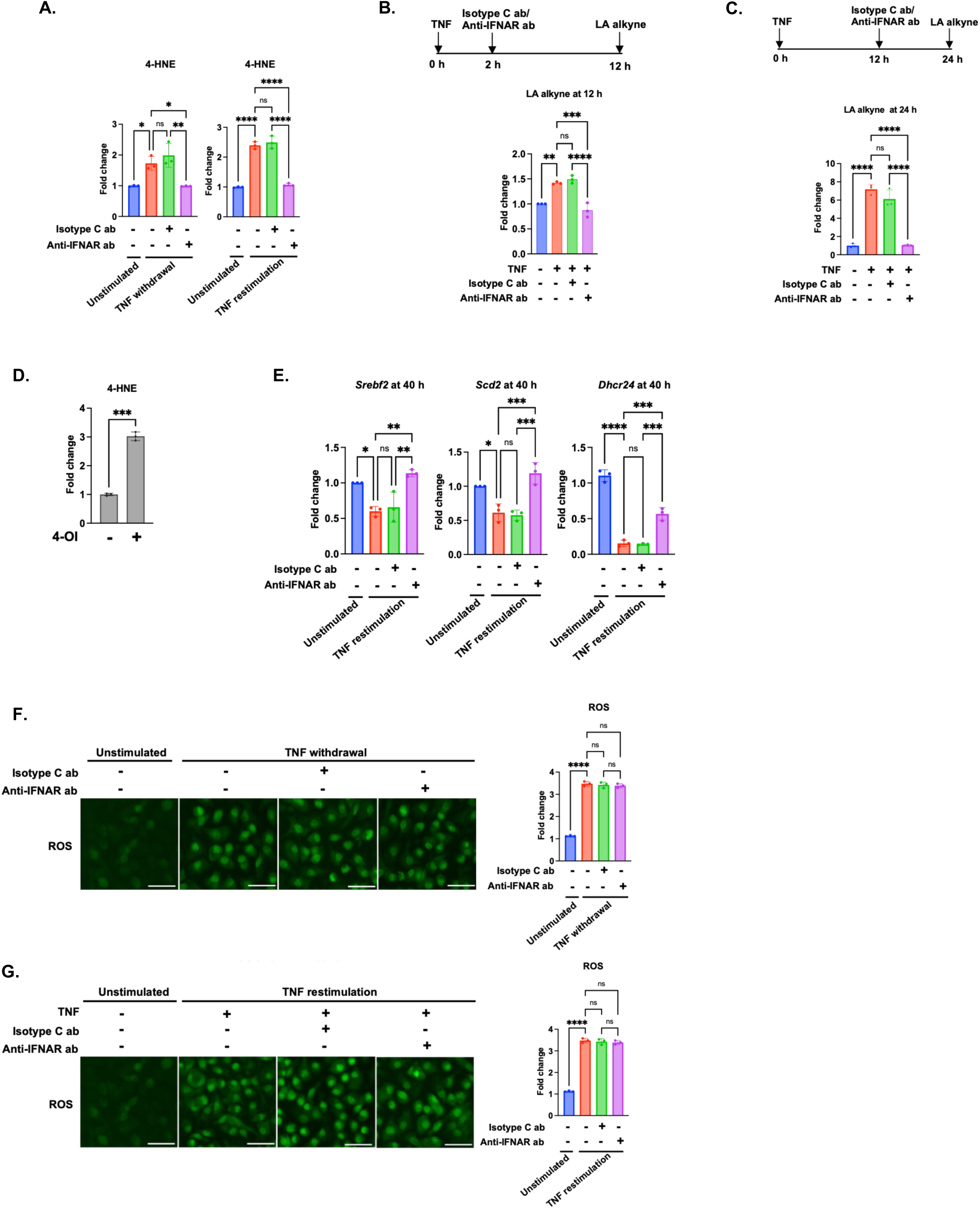
A. IFN-I sustains the 4-HNE adducts accumulation in B6.Sst1S BMDMs during pPAS. Bar graphs showing the fold change in fluorescence compared to the unstimulated samples. N=3. B and C. IFN-I initiates and maintains lipid peroxidation product synthesis in B6.Sst1S BMDMs during TNF stimulation. B6.Sst1S BMDMs were stimulated with TNF. Either at 2 h or 12 h, isotype control or anti-IFNAR antibodies were added to the culture, and the LA alkyne staining was carried out at either 12 h (B) or 24 h (C), respectively. Bar graphs showing the fold change in fluorescence compared to the respective untreated samples. N=3. D. Direct 4-Octyl itaconate treatment leads to 4-HNE adducts accumulation. B6.Sst1S BMDMs were treated with 4-OI for 24 h and the cells were immuno-stained for 4-HNE adducts. Bar graphs showing the fold change in fluorescence compared to the untreated samples. n=3. E. IFN-I downregulates the lipid biosynthesis gene expression during pPAS. Cells were harvested at 40 h for TNF restimulation condition and the mRNA levels of *Srebf2*, *Scd2*, and *Dhcr24* were quantified by RT-PCR. The fold change was calculated normalizing with the untreated control using ΔΔCt method using 18S as endogenous control. F and G. IFN-I do not regulate ROS during the pPAS. Confocal images showing the ROS during TNF withdrawal (F) or TNF restimulation (G) upon blocking IFN-I signaling. Scale bar-20 *μ*m. Bar graphs showing the fold change in fluorescence compared to the unstimulated samples. n=3. The data are presented as mean ± SD from three-five samples per experiment, representative of three independent experiments. The statistical analysis was performed by two-way ANOVA with Tukey’s multiple comparisons test (Panel A, E, F, and G), and one-way ANOVA with Tukey’s multiple comparisons test (Panel B and C). Significant differences are indicated with asterisks (ns, non significant; *, p < 0.05; **, p < 0.01; ***, p < 0.001; ****, p < 0.0001).

**Figure 3–figure supplement 1:**
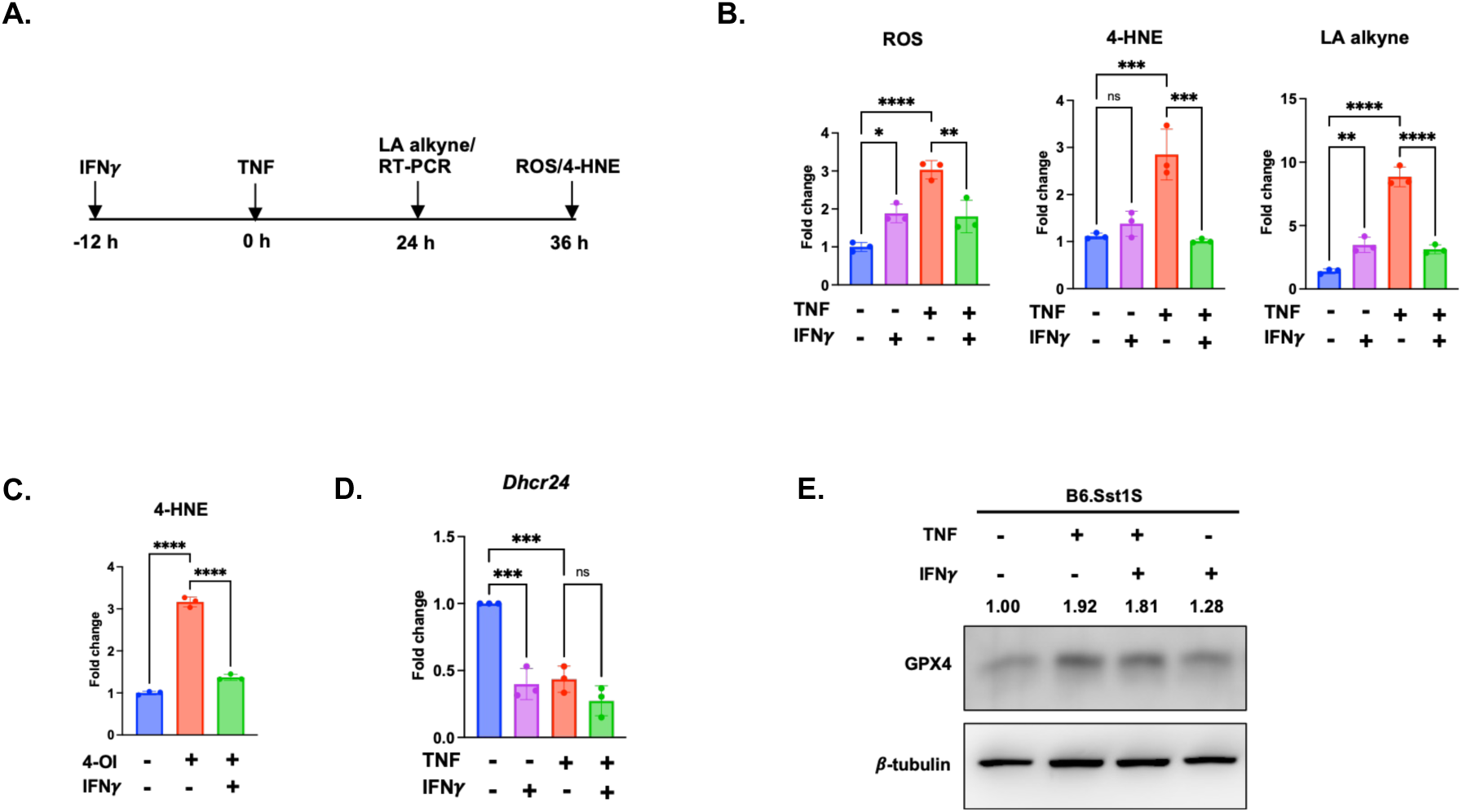
A. Experimental workflow. B6.Sst1S BMDMs were primed with IFN*γ* 12 h prior TNF stimulation. Cells were harvested either at 24 h or 36 h to understand the role of IFN*γ* on the IFN-I maintained pathological activation state. B. IFN*γ* priming reduces ROS levels during TNF stimulation. Upon IFN*γ* priming followed by TNF stimulation, B6.Sst1S BMDMs were stained with CellROX Green at 36 h. Bar graphs showing the fold change in fluorescence compared to the untreated samples. N=3. IFN*γ* priming prevents 4-HNE adducts accumulation during TNF stimulation. Upon IFN*γ* priming followed by TNF stimulation, B6.Sst1S BMDMs were immuno-stained for 4-HNE adducts at 36 h. Bar graphs showing the fold change in fluorescence compared to the untreated samples. N=3. IFN*γ* priming prevents lipid peroxidation product synthesis during TNF stimulation. Upon IFN*γ* priming followed by TNF stimulation, B6.Sst1S BMDMs were stained for LA alkyne at 24 h. Bar graphs showing the fold change in fluorescence compared to the untreated samples. n=3. C. IFN*γ* priming prevents 4-OI induced 4-HNE adducts accumulation. B6.Sst1S BMDMs were treated with 4-OI with or without IFN*γ* priming. Cells were immuno-stained for 4-HNE adducts at 36 h. Bar graphs showing the fold change in fluorescence compared to the untreated samples. n=3. D. IFN*γ* priming do not restore lipid biosynthesis gene expression during TNF stimulation. Upon IFN*γ* priming followed by TNF stimulation, B6.Sst1S BMDMs were harvested at 24 h and the mRNA levels of *Dhcr24* were quantified by RT-PCR. The fold change was calculated normalizing with the untreated control using ΔΔCt method using 18S as endogenous control. E. IFN*γ* priming do not enhance GPX4 expression. Upon IFN*γ* priming followed by TNF stimulation, B6.Sst1S BMDMs were harvested at 24 h GPX4 protein levels were quantified. Western blot images showing GPX4 protein expression considering *β*-tubulin as loading control. The average fold change in expression was mentioned compared to the untreated sample, normalized with *β*-tubulin levels. The data are presented as mean ± SD from three-five samples per experiment, representative of three independent experiments. The statistical analysis was performed by one-way ANOVA with Tukey’s multiple comparisons test (Panel B, C, and D). Significant differences are indicated with asterisks (ns, non significant; *, p < 0.05; **, p < 0.01; ***, p < 0.001; ****, p < 0.0001).

**Figure 4–figure supplement 1:**
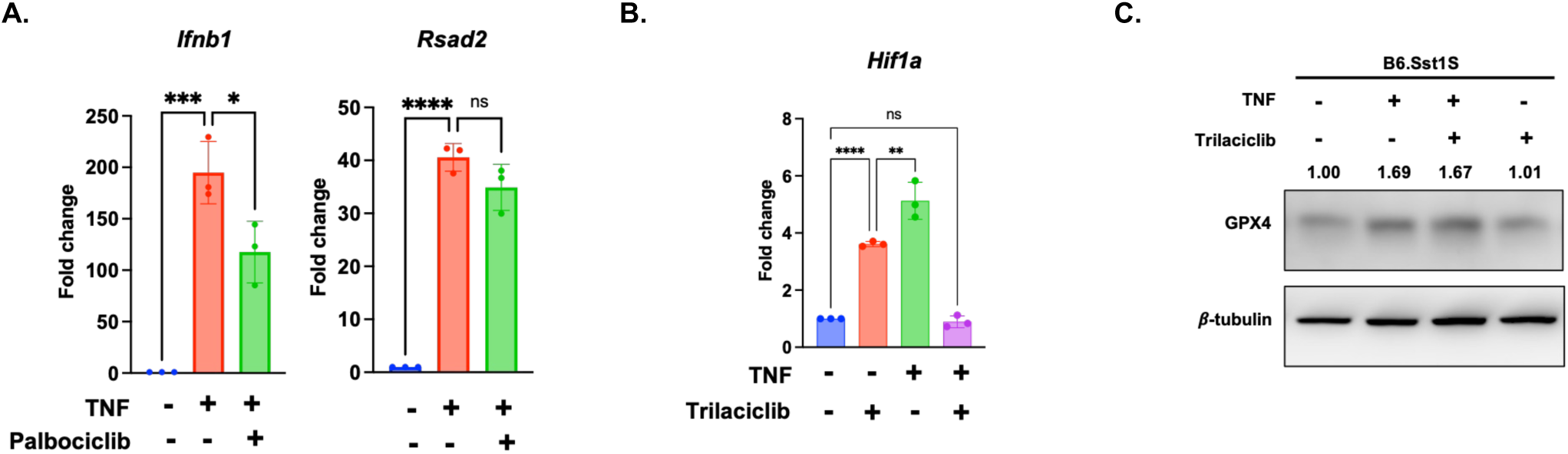
A. Palbociclib pretreatment partially prevents IFN-I super-induction during TNF stimulation. B6.Sst1S BMDMs were pretreated with palbociclib (3 *μ*M) for 12 h then with TNF. After 24 h of TNF stimulation, cells were harvested and the mRNA levels of *Ifnb1* and *Rsad2* were quantified by RT-PCR. The fold change was calculated normalizing with the untreated control using ΔΔCt method using 18S as endogenous control. B. Trilaciclib pretreatment enhances Hi1f1a during TNF stimulation. B6.Sst1S BMDMs were pretreated with trilaciclib (3 *μ*M) for 12 h then with TNF. After 24 h of TNF stimulation, cells were harvested and the mRNA levels of *Hif1a* were quantified by RT-PCR. The fold change was calculated normalizing with the untreated control using ΔΔCt method using 18S as endogenous control. C. Trilaciclib pretreatment do not enhance GPX4 expression. Upon Trilaciclib (3 *μ*M) pretreatment followed by TNF stimulation, B6.Sst1S BMDMs were harvested at 24 h GPX4 protein levels were quantified. Western blot images showing GPX4 protein expression considering *β*-tubulin as loading control. The average fold change in expression was mentioned compared to the untreated sample, normalized with *β*-tubulin levels. The data are presented as mean ± SD from three-five samples per experiment, representative of three independent experiments. The statistical analysis was performed by one-way ANOVA with Tukey’s multiple comparisons test (Panel A and B). Significant differences are indicated with asterisks (ns, non significant; *, p < 0.05; **, p < 0.01; ***, p < 0.001; ****, p < 0.0001).

**Figure 5–figure supplement 1:**
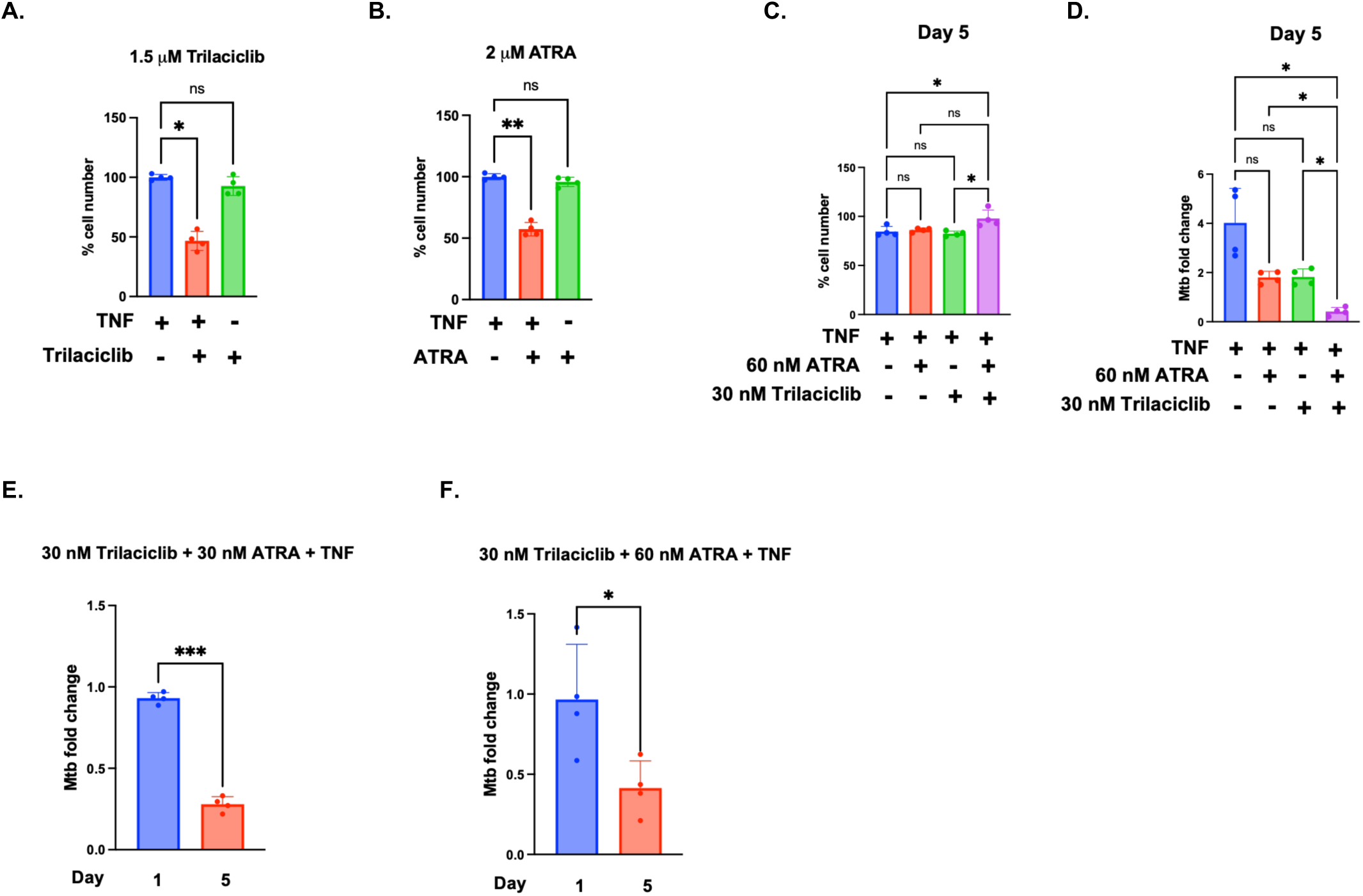
A - B. Higher concentration of Trilaciclib (1.5 *μ*M) and ATRA (2 *μ*M) induces cell loss in B6.Sst1S BMDMs under TNF stimulated and Mtb infected condition. B6.Sst1S BMDMs were pretreated with Trilaciclib (1.5 *μ*M) and ATRA (2 *μ*M) for 12 h followed by TNF stimulation for additional 24 h. Following day cells were infected with Mtb (MOI=1). Percent cell number was observed 5 days post infection using automated microscopy. (n=4). C – F. B6.Sst1S BMDMs were pretreated with trilaciclib (30 nM) and ATRA (30 nM or 60 nM), either alone or in combination, for 12 h followed by TNF stimulation (10 ng/mL) for an additional 24 h. Cells were subsequently infected with the Mtb Erdman reporter strain (SSB-GFP, smyc’::mCherry) at an MOI of 1 for 5 days. Total cell numbers were quantified by automated microscopy on days 1 and 5 post-infection (C). Combination treatment with 30 nM trilaciclib and either 30 nM or 60 nM ATRA reduced intracellular Mtb growth in B6.Sst1S BMDMs, as determined by qPCR-based assay on day 5 post-infection (D – F). Bar graphs show the fold change in Mtb burden from day 1 to day 5 post-infection following combination treatment. Data represent mean ± SEM from four independent experiments (n=4). The data are presented as mean ± standard deviation (SD) from three-five samples per experiment, representative of three independent experiments. The statistical analysis was one-way ANOVA with Tukey’s multiple comparisons test. Significant differences are indicated with asterisks (ns, non significant; *, p < 0.05; **, p < 0.01; ***, p < 0.001; ****, p < 0.0001).

