## Supplementary material for "IFN*γ* and IFN*γ* mimetics prevent IFN-I-mediated TB susceptibility by regulating iron metabolism and lipid peroxidation": Key Resources Table

| **Key Resources Table** | | | | |
| --- | --- | --- | --- | --- |
| **Reagent type (species) or resource** | **Designation** | **Source or reference** | **Identifiers** | **Additional information** |
| Antibody | Mouse monoclonal anti-IFNAR1 antibody (clone MAR1-5A3), functional grade | Thermo Fisher Scientific | Cat# 16594585, RRID:AB_1210688 | IFN I inhibition (1:200) |
| Antibody | Mouse IgG1 κ isotype control antibody (clone P3.6.2.8.1) | Thermo Fisher Scientific | Cat# 14471482, RRID:AB_470111 | Isotype C Ab (1:200) |
| Antibody | Mouse monoclonal anti-TNFα antibody (clone XT22) | Thermo Fisher Scientific | Cat# MM350D, RRID:AB_223528 | Inhibition of TNF signalling (10 mg/mL) |
| Antibody | Mouse monoclonal anti-β-tubulin antibody | Santa Cruz Biotechnology | Cat# sc-55529, RRID:AB_2210962 | WB (1:1000) |
| Antibody | Rabbit monoclonal anti-Ferritin Heavy Chain antibody [EPR18878] | Abcam | Cat# ab183781, RRID:AB_2940987 | WB (1:2000) |
| Antibody | Rabbit monoclonal anti-Ferritin Light Chain antibody [EPR5260] | Abcam | Cat# ab109373, RRID:AB_1086271 | WB (1:5000) |
| Antibody | Rabbit polyclonal anti-4-Hydroxynonenal antibody | Abcam | Cat# ab46545, RRID:AB_722490 | IF (1:100) |
| Antibody | Mouse monoclonal anti-GPX4 antibody | Proteintech Group Inc | Cat# 67763-1-Ig, RRID:AB_2909469 | WB (1:1000) |
| Antibody | HIF-1α (D2U3T) Rabbit monoclonal antibody | Cell Signaling Technology | Cat#  14179S  RRID:AB_2622225 | WB (1:1000) |
| Antibody | Horse anti-mouse IgG, HRP-linked antibody (polyclonal) | Cell Signaling Technology | Cat# 7076s, RRID:AB_330924 | WB (1:2000) |
| Antibody | Goat anti-rabbit IgG, HRP-linked antibody (polyclonal) | Cell Signaling Technology | Cat# 7074s, RRID:AB_2099233 | WB (1:2000) |
| Antibody | F(ab')2-Goat anti-Rabbit IgG (H+L) Cross-Adsorbed Secondary Antibody, Alexa Fluor™ 488, Invitrogen™ | Thermo Fisher | Cat#  A11070  RRID:AB_2534114 | IF (1:500) |
| Strain, strain background (*Mycobacterium bovis* BCG*)* | *Mycobacterium bovis* BCG, TMC 1019 [BCG Japanese] | ATCC | Cat# 35737 |  |
| Strain, strain background (*Mycobacterium tuberculosis)* | Erdman (SSB-GFP, *smyc′*::mCherry) | (Lavin & Tan, 2022) | N/A | A gift from Shumin Tan |
| chemical compound, drug, peptides | Deferriprone | Thermo Fisher | Cat#  501363627 |  |
| chemical compound, drug, peptides | Murine IFN-gamma | Peprotech | Cat# 315-05 |  |
| chemical compound, drug, peptides | Murine TNF-alpha | Peprotech | Cat# 315-01A |  |
| chemical compound, drug, peptides | Mouse IFN-Beta, Mammalian | Thermo Scientific | Cat#  50-153-7986 |  |
| chemical compound, drug, peptides | Middlebrook 7H9 Broth | BD Biosciences | Cat# 271310 |  |
| chemical compound, drug, peptides | Middlebrook 7H10 Agar | BD Biosciences | Cat# 262710 |  |
| commercial assay or kit | TaqMan™ Environmental Master Mix 2.0 | Fisher Scientific | Cat#4396838-5mL |  |
| commercial assay or kit | CellROX™ Green Reagent | Thermo Fisher Scientific | Cat# C10444 |  |
| commercial assay or kit | FerroOrange | Cell Signaling Technology | Cat#36104S |  |
| commercial assay or kit | Click-iT™ Lipid Peroxidation Imaging Kit | Thermo Fisher Scientific | Cat# C10446 |  |
| commercial assay or kit | RNeasy plus mini kit | Qiagen | Cat#74136 |  |
| commercial assay or kit | TRIzol Reagent | Invitrogen | Cat#15596026 |  |
| commercial assay or kit | PrimeScript™ RT Master Mix (Perfect Real Time) | Takara Bio USA | Cat#1  RR036A |  |
| commercial assay or kit | GoTaq qPCR Mastermix | Promega | Cat#A6002 |  |
| Strain, strain background (*Mus musculus)* | Mouse: C57BL/6J, adult male and female | The Jackson Laboratory | Stock No.: 000664, RRID:IMSR_JAX:000664 | <https://www.jax.org/strain/000664> |
| Strain, strain background (*Mus musculus)* | Mouse: B6J.C3-*Sst1^C3HeB/Fej^*Krmn, adult male and female | Pichugin et al, 2009. | Stock No: 043908-UNC  <https://www.mmrrc.org> | Available at <https://www.mmrrc.org> |
| Strain, strain background (*Mus musculus)* | Mouse: (C3XB6.Sst1S) F1, adult male | Yabaji et al, 2025. | N/A |  |
| sequence-based reagent | Mtb specific_F | Integrated DNA Technologies, Inc. | PCR primers | GGAAATGTCACGTCCATTCATTC |
| sequence-based reagent | Mtb specific_R | Integrated DNA Technologies, Inc. | PCR primers | GCGTTGTTCAGCTCGGTA |
| sequence-based reagent | Mtb specific probe | Integrated DNA Technologies, Inc. | PCR probe | 56-FAM/AGCTTGGTCAGGGACTGCTTCC/36-TAMSp/ |
| sequence-based reagent | BCG specific_F | Integrated DNA Technologies, Inc. | PCR primers | GTGGTGGAGCGGATTTGA |
| sequence-based reagent | BCG specific_R | Integrated DNA Technologies, Inc. | PCR primers | CAACCGGACGGTGATCC |
| sequence-based reagent | BCG specific probe | Integrated DNA Technologies, Inc. | PCR probe | /5Cy5/TTCTGGTCG/TAO/ACGATTGGCACATCC/3IAbRQSp/ |
| nucleotide sequence | *Ciita_*F | Integrated DNA Technologies, Inc. | PCR primers | CTTCAAGCAGCCTCAGTATC |
| nucleotide sequence | *Ciita_*R | Integrated DNA Technologies, Inc. | PCR primers | ATGTGTCCTCTGTCTCATTTAC |
| nucleotide sequence | *Ifnb1_*F | Integrated DNA Technologies, Inc. | PCR primers | ATGAGTGGTGGTTGCAGGC |
| nucleotide sequence | *Ifnb1_*R | Integrated DNA Technologies, Inc. | PCR primers | TGACCTTTCAAATGCAGTAGATTC |
| nucleotide sequence | *Rsad2_*F | Integrated DNA Technologies, Inc. | PCR primers | AAGCTGAGGAGGTGGTGCAG |
| nucleotide sequence | *Rsad2_*R | Integrated DNA Technologies, Inc. | PCR primers | GAAAACCTTCCAGCGCACAG |
| nucleotide sequence | *Srebf2_F* | Integrated DNA Technologies, Inc. | PCR primers | GGACAGTGATGTGGACTTGAA |
| nucleotide sequence | *Srebf2_R* | Integrated DNA Technologies, Inc. | PCR primers | CGGCTCAGAGTCAATGGAATAG |
| nucleotide sequence | *Scd2_F* | Integrated DNA Technologies, Inc. | PCR primers | AGCACCTTCTTGCGATACG |
| nucleotide sequence | *Scd2_R* | Integrated DNA Technologies, Inc. | PCR primers | GATGTTCTCCCGAGAGCTAATG |
| nucleotide sequence | *Dhcr24_F* | Integrated DNA Technologies, Inc. | PCR primers | GAGTCATCGTCCCACAAGTTATG |
| nucleotide sequence | *Dhcr24_R* | Integrated DNA Technologies, Inc. | PCR primers | GGCATAGAACAGGTCTGAGTTT |
| nucleotide sequence | *Hif1a_F* | Integrated DNA Technologies, Inc. | PCR primers | CCCATTCCTCATCCGTCAAATA |
| nucleotide sequence | *Hif1a_R* | Integrated DNA Technologies, Inc. | PCR primers | GGCTCATAACCCATCAACTCA |
| nucleotide sequence | *Acod1_F* | Integrated DNA Technologies, Inc. | PCR primers | GTATCATTCGGAGGAGCAAGAG |
| nucleotide sequence | *Acod1_R* | Integrated DNA Technologies, Inc. | PCR primers | GGAGGTGTTGGAACTGTAGATT |
| sequence-based reagent | *Trib3_*F | Integrated DNA Technologies, Inc. | PCR primers | GCAAAGCGGCTGATGTCTG |
| sequence-based reagent | *Trib3_*R | Integrated DNA Technologies, Inc. | PCR primers | AGAGTCGTGGAATGGGTATCTG |
| sequence-based reagent | *Chac1_*F | Integrated DNA Technologies, Inc. | PCR primers | CCTGCTACCCTGCTCTTACCT |
| sequence-based reagent | *Chac1_*R | Integrated DNA Technologies, Inc. | PCR primers | GAGCTTGGCTCCTCAGGTC |
| sequence-based reagent | *b-actin_*F | Integrated DNA Technologies, Inc. | PCR primers | GTGGGCCGCTCTAGGCACCA |
| sequence-based reagent | *b-actin_*R | Integrated DNA Technologies, Inc. | PCR primers | CGGTTGGCCTTAGGGTTCAGGG |
| sequence-based reagent | *18S_*F | Integrated DNA Technologies, Inc. | PCR primers | TCAAGAACGAAAGTCGGAGGT |
| sequence-based reagent | *18S_*R | Integrated DNA Technologies, Inc. | PCR primers | CGGGTCATGGGAATAACG |
| software, algorithm | GraphPad Prism V 10.6.1 | GraphPad Software, LLC. | <https://www.graphpad.com/>, RRID: SCR_002798 |  |
| software, algorithm | Microsoft office | Microsoft | https://www.office.com/?auth=2 |  |
| software, algorithm | EndnoteX9 | Clarivate Analytics | https://endnote.com/downloads |  |
| software, algorithm | ImageJ | National Institutes of Health (NIH) | <https://imagej.nih.gov/ij/>, SCR_003070 | Image analysis |
| software, algorithm | GSEA | GSEA | RRID: SCR_003199 |  |
| software, algorithm | Seurat | Seurat | RRID: SCR_007322 |  |
| software, algorithm | RStudio | RStudio | RRID: SCR_000432 |  |
| other | Operetta CLS HCA System | Operetta^TM^ | https://www.perkinelmer.com/in/lab-solutions/product/operetta-cls-system-hh16000020 |  |
| other | SP5 Confocal Microscope | Leica | N/A |  |
| other | LAS-4000 | FujiFilm | N/A |  |
| other | ProLong™ Gold Antifade Mountant | Invitrogen™ | Cat# P36934 |  |
| other | Hoechst 33342 | Fisher Scientific | Cat# H3570 | 10 mg/mL |
| other | Paraformaldehyde Solution 4% in PBS | Fisher Scientific | Cat# J19943-K2 |  |
| other | L-Glutamine | Corning® | Cat# 25-005-CI |  |
| other | Penicillin Streptomycin solution | Corning® | Cat# 30-002-CI |  |
| other | HEPES buffer | Corning® | Cat# 25-060-CI |  |
| other | L929 Cell Conditioned Media (LCCM) | This paper | N/A |  |
| other | Lymphoprep^TM^ (1.077A) | STEMCELL | Cat#07801 |  |
| other | Poly Ethylene Glycol (PEG), Bioultra-8000 | Sigma | Cat#89510 |  |
| other | 5M NaCl | Invitrogen | Cat#AM9759 |  |
| other | Tris Hydrochloride, 1M solutions (pH 8.0) | Fisher Scientific | Cat#77-86-1 |  |
| other | Ultrapure 0.5 M EDTA pH 8.0 | Invitrogen | Cat#15575-038 |  |
| other | Ambion^TM^ Nuclease-free Water | Invitrogen | Cat#AM9932 |  |
| other | SpeedBead Magnetic Carboxylate Modified Particles | GE Healthcare | Cat#65152105050250 |  |
| other | DynaMag^TM^-96 side | Life Technologies^TM^ | Cat#12331D |  |
| other | Glycine | Sigma | Cat#50046 |  |
| other | NaOH Solution | Sigma | Cat#72068 |  |
| other | Proteinase K | Ambion | Cat#AM2546 |  |
| chemical compound | Trilaciclib dihydrochloride | Selleckchem | Cat#  E4578 |  |
| chemical compound | Retinoic acid/ATRA | Selleckchem | Cat#  S1653 |  |
